# Early human fetal lung atlas reveals the temporal dynamics of epithelial cell plasticity

**DOI:** 10.1101/2023.10.27.564403

**Authors:** Henry Quach, Spencer Farrell, Kayshani Kanagarajah, Michael Wu, Xiaoqiao Xu, Prajkta Kallurkar, Andrei Turinsky, Christine E. Bear, Felix Ratjen, Sidhartha Goyal, Theo J. Moraes, Amy P. Wong

**Affiliations:** Program in Developmental and Stem Cell Biology, Hospital for Sick Children, 686 Bay Street, PGCRL 16-9420, Toronto, Ontario M5G0A4; Department of Laboratory Medicine & Pathobiology, University of Toronto; Department of Physics, University of Toronto; Centre for Computational Medicine, Hospital for Sick Children, Toronto, Canada; Program in Molecular Medicine, Hospital for Sick Children, Toronto, Canada; Program in Translational Medicine, Hospital for Sick Children, Toronto, Canada

**Keywords:** Lung development, Single-cell RNA and spatial transcriptomics, pluripotent stem cell differentiation, human fetal lung

## Abstract

While animal models have provided key insights into conserved mechanisms of how the lung forms during development, human-specific developmental mechanisms are not always captured. To fully appreciate how developmental defects and disease states alter the function of the lungs, studies in human lung models are important. Here, we sequenced >150,000 single single-cells from 19 healthy human fetal lung tissues from gestational weeks 10-19 and identified at least 58 unique cell types/states contributing to the developing lung. We captured novel dynamic developmental trajectories from various progenitor cells that give rise to club, ciliated, and pulmonary neuroendocrine cells. We also identified four CFTR-expressing progenitor cell types and pinpointed the temporal emergence of these cell types. These developmental dynamics reveal broader epithelial cell plasticity and novel lineage hierarchies that were not previously reported. Combined with spatial transcriptomics, we identified both cell autonomous and non-cell autonomous signalling pathways that may dictate the temporal and spatial emergence of cell lineages. Finally, we showed that human pluripotent stem cell-derived fetal lung models capture cell lineage trajectories specifically through CFTR-expressing progenitor cells, that were also observed in the native fetal tissue. Overall, this study provides a comprehensive single-cell atlas of the developing human lung, outlining the temporal and spatial complexities of cell lineage development.

**Highlights:**

1. Single-cell transcriptomics atlas from 19 human fetal lungs reveals cellular heterogeneity and previously unappreciated cellular plasticity in the epithelial compartment.
2. Identification of novel CFTR-expressing progenitor cells that gives rise to club, ciliated and PNEC.
3. Novel RNA velocity facilitated the identification of dynamic lineage trajectories in the epithelial compartment.
4. Temporally regulated cell signaling through promiscuous interactions between sender and receiving cells may dictate cell lineage fates.
5. Integration of human pluripotent stem cell (hPSC)-derived fetal lung cells and organoids with primary lung dataset show hPSC-differentiations captures key developmental trajectories of fetal epithelial cell states.

## Introduction

Single-cell technologies are redefining the molecular signatures of canonical cell types and identifying new cell types and cell states in organs and tissues^1,2^. Specifically, single-cell RNA sequencing (scRNA-seq) has revealed cellular transcriptomic heterogeneities and has enriched our understanding of how these cells contribute to development and disease. Single-cell atlases of the adult human pulmonary system have illuminated the roles of key cell types in homeostasis and some diseases^3–6^. On the contrary, our understanding of the cells that form the developing fetal lung has largely been extrapolated from mouse studies^7,8^, and not until recently, have there been studies leveraging scRNA-seq to characterize the developing human fetal lung tissue^9,10^. Through single cell studies, species-specific differences in lung cell types and cellular distributions have been identified, which can impact function and disease pathogenesis. Understanding the molecular and cellular networks that drive the formation of the respiratory system during fetal development may provide important insight into the regenerative processes during repair and potentially identify developmental origins of postnatal lung disease/disorders.

Recent sequencing datasets from a small collection of normal human fetal lungs obtained from early to mid-gestation fetuses, have highlighted some novel cell types and cell lineage trajectories^9,11^ that are unique to human fetal development. Unfortunately, following up on these observations and their potential impact in post-natal lung function is hindered by lack of experimentally tractable models.

Human pluripotent stem cells (hPSCs) are becoming a resourceful tool to generate “organoids” or in-vitro tissue mimetics to study fundamental mechanisms of human cell lineage development. Directed differentiation protocols towards mature lung epithelia^12–14^ aim to recapitulate stepwise developmental milestones to ensure proper generation of specific cells and to date has achieved robust differentiations that are enabling cell therapy discoveries for the treatment of lung diseases such as Cystic fibrosis^15–17^. However, to precisely model the native tissue, a comprehensive understanding of the cells that exist and how they develop requires in-depth analysis of the native tissue. Understanding the developmental trajectories and pathways driving these cell lineage formations will also improve current hPSC differentiation protocols to generate bona fide cell types.

Here, we present a comprehensive single-cell atlas of freshly isolated human fetal lungs from 19 independent lung samples spanning gestational weeks 10-19 and capturing >150,000 cells in total. Based on differentially expressed genes (DEG), we identified 58 cell types in the developing fetal lungs from the integrated datasets of all 19 lung samples. While the presence of canonical cell types previously found in both adult human lungs^4^ and embryonic mouse lungs^8^ were observed, we also identify novel cell types that are uniquely found in human fetal lungs including CFTR-expressing epithelial progenitor cells, a *PTEN+STAT3+* distal epithelial progenitor, and an uncommitted stromal cell-like epithelial cell. Given the large number of cells sequenced from our collection of fetal tissues, our dataset reveals important dynamic epithelial cell plasticity and new lineage trajectories not previously identified. To determine cell lineage relationships and the ancestral origins of specific cell types, we leveraged our previously established modified RNA velocity prediction tool, LatentVelo^18^, followed by partition-based graph abstraction (PAGA) visualization and *Slingshot*^19^ to infer novel trajectories and show the cellular transitions of the cells from different cell origins. We identified classical trajectories previously found in the immune, stromal and endothelial compartment, but we also uncovered novel unexpected trajectories in the developing epithelium. We revealed the dynamic nature of how CFTR-expressing epithelial progenitors contribute to the pulmonary neuroendocrine and multi-ciliated lineages. Moreover, in combination with spatial transcriptomics, we revealed cell dynamic nature of signalling pathways via sender and receiver cells that may contribute to cell fates changes, specifically through the *SCGB3A2+SFTPB+CFTR+* cell population. Finally, we show that hPSC-derived fetal lung epithelia models capture several unique epithelial cell progenitors and lineage trajectories observed in the native fetal airways and supports the use of hPSC-derived fetal lung models for future studies in specific cell lineage development and function during development.

## Results

### Cell composition of the human fetal lungs

A total of 19 freshly isolated fetal lung tissues from elective pregnancy terminations from gestational weeks (GW)10-19 were collected and processed for 3’ single-cell RNA sequencing (scRNA-seq) using from 10X Genomics. One sample was sampled twice from different regions of the lung to capture more cells representing from the upper and lower lungs (denotated with an asterisk, **Supplementary Table 1**). A total of 170,256 single-cells were sequenced at approximately 60,000 read depth per cell. FASTQ files were generated, and the reads were aligned with human reference genome hg38 (GRCh38) (**Extended Data Fig. 1A**). After the removal of low-quality cells, dead cells, and doublets, 156,698 cells were analyzed. After normalization and dimensionality reduction, reciprocal principal component analysis (rPCA) was used to identify anchors for integration to generate the fetal lung dataset. Based on the expression of the male sex-determining genes*, SRY and DDX3Y*, and the X-chromosome inactivation gene, *XIST*, 12 male and 7 female lungs were collected (**Fig. 1A**). Leiden clustering informed by *Clustree*^20^ was used to unbiasedly define the number of cell clusters (**Extended Data Fig. 1B**). Differentially expressed genes (DEG) based on non-parametric Wilcoxon rank sum test were used to annotate the cell populations which were broadly classified into 5 uniquely different cell populations (**Fig. 1B** and **Supplementary Table 2**) designated as stromal (N = 98,166; *COL1A1, COL1A2*), epithelial (N = 16,068; *EPCAM*), endothelial (N = 13,376; *PECAM1*), immune (N = 16,258; *CD74, CD3, NKG7*), and Schwann cells (N = 512; *NRXN1, S100B, PLP1*) (**Fig. 1C**). Further sub-clustering of each main cell population was informed by *Clustree* and DEG which resulted in a total of 58 distinct cell types/states in the developing lungs (**Fig. 1D**). Assessing the cell proportions over gestational time showed stromal cells formed the largest proportion of cells throughout the 10 weeks of development (**Extended Data Fig. 1C**). Temporal assessment based on the gestational week of the lung tissues highlights the dynamic changes of cell populations (immune cells, red hatched box), with emergence of cells (ciliated cells, black hatched box), and temporal disappearance (specific epithelial subpopulations, green hatched box) of cell types over time (**Fig. 1E**). Cell type proportions of these cell types showed changes in the presence of varying cell types **(Supplementary Table 3).**

**Fig 1:**
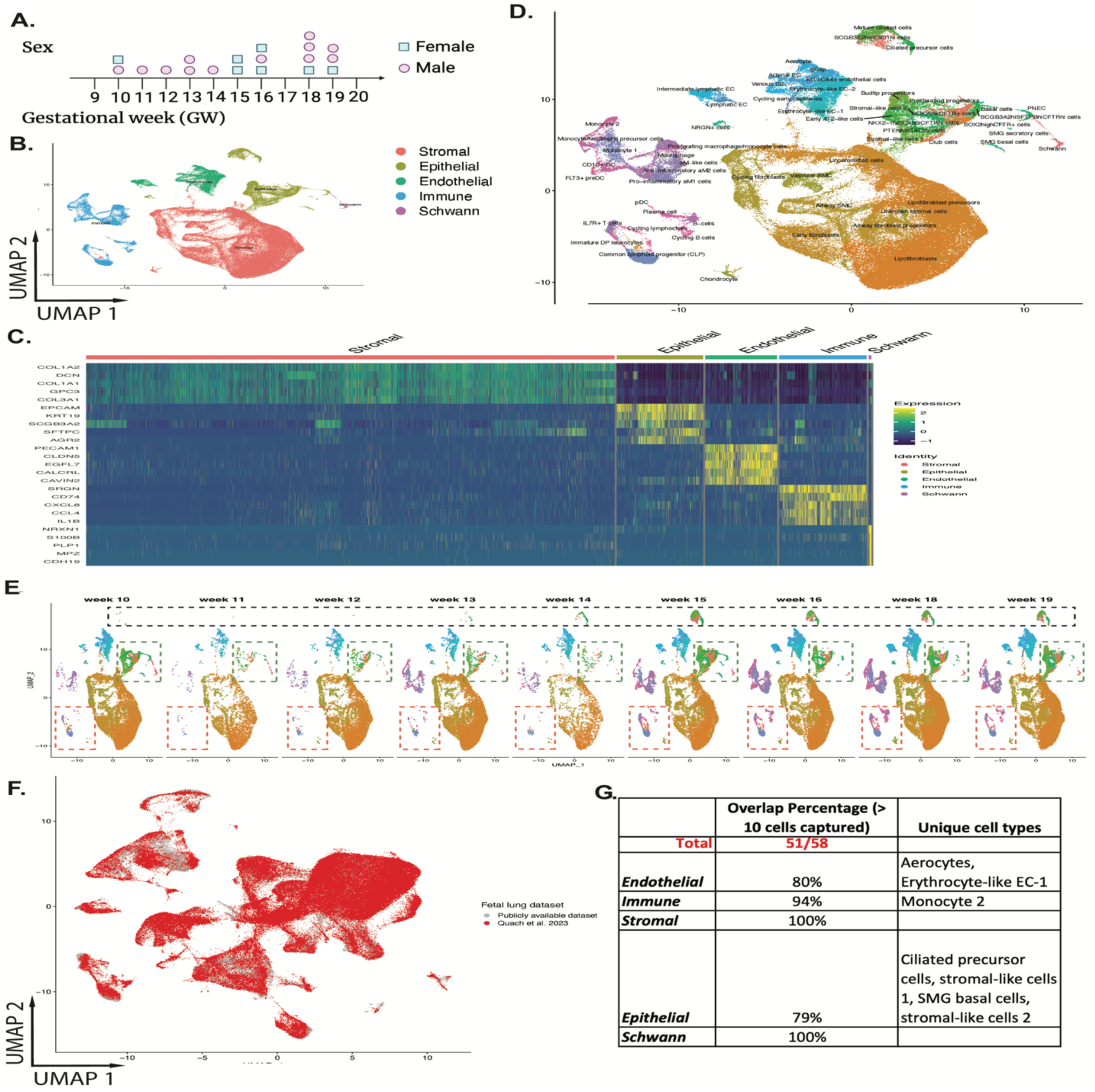
Overview of the single-cell RNA sequencing dataset from 19 fetal lung tissues. **A:** Biological sex of the lungs was determined by *SRY, XIST*, and *DDX3Y* expression. **B:** UMAP visualization of the dataset highlighting the main cell types: stromal, epithelial, endothelial, immune, and schwann cells. **C:** Gene expression heatmap of the top 5 differentially expressed genes in each cluster. **D:** UMAP projection of all 58 cell types/states identified within the integrated dataset. The number of clusters were determined by *Clustree*. **E:** UMAP projection of the integrated fetal lung datasets annotated by gestational week (GW). **F:** UMAP projection of two recently published fetal lung datasets (He et al. and Sountoulidis et al.) overlayed onto our dataset as determined by single-cell reference mapping (Seurat). **G:** Proportion of cell types shared and unique to our dataset.

We performed an integrated analyses to identify similarities and differences between our dataset and two recently published datasets encompassing similar developmental lung tissues^9,10^. Integrated UMAP overlay of our dataset (in red, **Fig. 1F**) with the published two datasets by He et al.^9^, and Sountoulidis et al.^10^, (combined, in grey) showed that 51 of the 58 cell clusters (∼88%) were common amongst all three datasets (**Fig. 1G** and **Extended Data Fig. 1D**). However, 7 unique cell types were exclusively captured in our dataset and these included aerocytes and erythrocyte-like cell 1 from the endothelial cluster, monocyte 2 from the immune cluster, and stromal-like 1, stromal-like 2, ciliated cell precursor, and SMG basal cell in the epithelial cluster.

This may reflect the larger cell number or the comprehensive range of fetal tissue that enabled us to capture these cell types/states. While He et al.^9^ identified “aerocytes” and SMG basal cells in their dataset, differences between He et al.’s and our dataset’s cell type classification was found. Interestingly, our dataset captured developing aerocytes (*TBX2, EDNRB, APLN*) that appeared as early as GW10, whereas He et al. only captured more mature aerocytes (TBX2, S100A3) at GW22. Differences in SMG basal cells can be accounted by inclusion of *CNN1, LGALS3,* and *LGALS1* genes to define these cells in our dataset^21^.

We next compared our fetal lung dataset to published single-cell atlas from human adult lung^4^ using *MapQuery* function in Seurat to map query cells onto our reference fetal lung dataset. While we only identified 47% cell types that were shared between our dataset and the adult lung dataset (**Extended Data Fig. 1E**), it is plausible that there are more significant common cell types shared that have not been captured due to the limited sample size of the adult lung dataset used. Cells that are common between fetal and adult lung tissues include many of the canonical cell types including endothelial subtypes, immune cells, and some epithelial subtypes (SMG secretory and basal cells, airway basal cells, and ciliated cells). As expected, unique fetal lung cell types were exclusively observed in our dataset (ie. Budtip progenitors, and several CFTR+ progenitor cells).

We also compared our dataset to a previous mouse embryonic lung single-cell atlas^8^ and found 45% shared cell types (**Extended Data Fig. 1F**). Notable similarities in the cells captured from the mouse dataset included several endothelial, immune and stromal subtypes. It should be noted that the mouse lung dataset by Negretti et al captures a broad range of tissues from embryonic day 12 (equivalent to the pseudoglandular lung tissues in mice). The large number of cells not captured in our fetal lung dataset that are found in the mouse embryonic lung dataset may be attributed to the differences in developmental stages of the tissue, which would capture different cell types/states. Other explanations include discordance in cell cluster annotation and differences in single-cell preparation, including MACS enrichment used in several single cell studies that would exclude specific cell types.

### The developing lung stroma

Stromal cells formed the largest proportion of cells in all lung tissues analyzed (**Fig. 1B**). We identified 10 stromal cell subtypes (**Fig. 2A**) based on DEG expression (**Supplementary Table 2**). These included lipofibroblasts, lipofibroblast precursors, cycling fibroblasts, early fibroblasts, airway fibroblast progenitors, uncommitted cells, airway smooth muscle cells (SMC), vascular SMC, chondrocytes, and an unknown stromal cell population. Lipofibroblasts and their precursors were the most abundant stromal cell type across all gestational weeks (**Fig. 2B**). Both cell types expressed abundant *TCF21*, previously shown in both mouse fetal and adult lung lipofibroblasts^22^. Lipofibroblast precursors differentially expressed higher levels of several canonical mesenchymal associated genes including the pleckstrin homology domain family H member 2 (*PLEKHH2),* and the microtubule actin cross-linking factor (*MACF1*) (**Fig. 2C**) as previously shown in mouse lipofibroblasts cell lineages^23^. Lipofibroblasts expressed progressively higher levels of the transcription factors *JUN, FOS* and *EGR1* (**Fig. 2D**), which are known to regulate the synthesis of structural extracellular matrices (ECM) such as collagen and elastin which are important in alveolar development^24^. While alveolar development occurs later in gestation, it is conceivable that lipofibroblasts in early fetal lung development play a role in laying down the ECM proteins required to form the airway structures. Differential enrichment analysis of the transcription factors driving these cell lineages showed an enrichment of *JUN*, *FOS,* and *IRF1* expression in lipofibroblasts and *STAT2* and *GATAD1* expression in lipofibroblast precursors.

**Fig 2:**
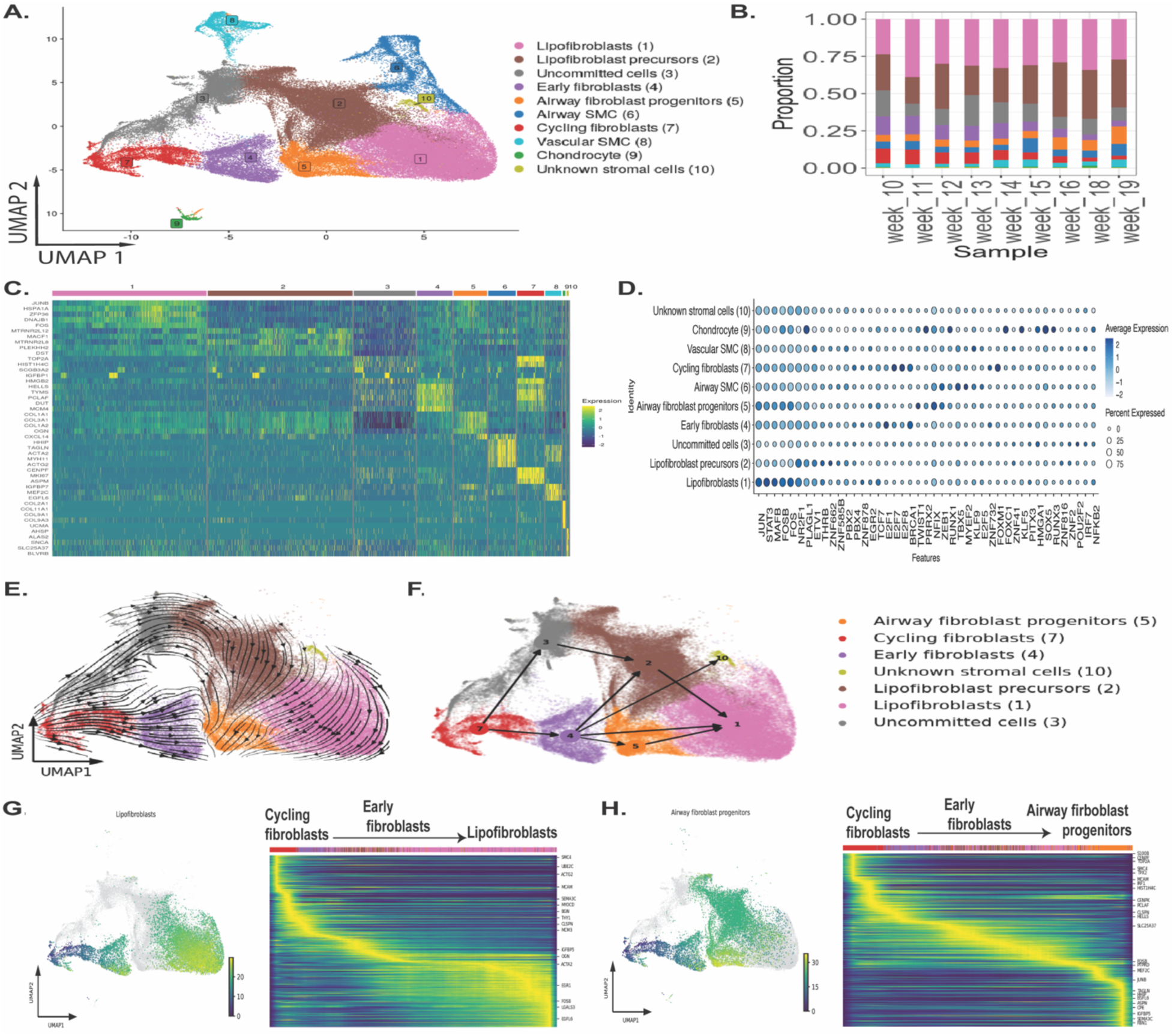
Characterization of the stromal cell compartment identifies lipofibroblast and lipofibroblast progenitors as main cell type in the developing fetal lungs. **A:** UMAP visualization of the fetal stromal cell subtypes. **B:** Proportion of stromal cell subtypes across gestational week. **C:** Gene expression heatmap of the top 5 differentially expressed genes in each cluster. **D:** Dotplot of the top differentially expressed transcription factors (TF) genes based on regulon specificity score (RSS) via *SCENIC*. **E:** LatentVelo analysis revealing inferred cell state trajectories projected onto the UMAP. **F:** Partition-based graph abstraction (PAGA) plot identified lineage trajectories informed by LatentVelo velocities. **G:** Slingshot trajectory analysis with a root at Cycling fibroblasts identifies a trajectory to Lipofiblasts. Trajectory heatmap plot shows the progression of cell types along the trajectory, and the change in significantly varying genes. **H:** Slingshot trajectory analysis with a root at Cycling fibroblasts identifies a trajectory to Airway fibroblast progenitors.

Cycling fibroblasts and early fibroblasts were also present at a relatively high proportion in the early lung tissues (GW 10-14; ∼6-10% and ∼9-12% respectively) but gradually reduced in later gestational lung tissues (GW15 onwards; ∼2-5% and ∼3-7% respectively) (**Fig. 2B, Supplementary Table 3**). Other than the common collagen genes (*COL1A1, COL1A2*), cycling fibroblasts expressed high levels of genes associated with cellular proliferation *TOP2A, CENPF, MKI67* (**Fig. 2C**). Early fibroblasts expressed high levels of genes associated with chromatin remodeling such as *HELLS* and *TYMS*^25^(**Fig. 2D**), previously shown to be highly expressed in mouse embryonic fibroblasts. The transcription factors differentially expressed in these cells include *E2F7, FOXM1* (known regulators of cell cycle in fibroblasts^26^) and *MYBL2*, a regulator of cell cycle progression, respectively. Interestingly, tight regulation of *MYBL2* has previously been shown to be critical in somatic reprogramming of mesenchymal to epithelial transitions^27^, suggesting a potential contribution of these stromal cells to the epithelium.

There was also a significant proportion of uncommitted cells that were closely associated with cycling fibroblasts and lipofibroblast precursors based on the UMAP projections (**Fig. 2A**). These uncommitted cells share some transcriptional similarities to cycling fibroblasts and lipofibroblast precursors (**Fig. 2C**), but uniquely expressed high levels of the tissue inhibitor of matrix metalloproteinases *(TIMP3)* and galectin-3 *(LGALS3),* both involved in regulating the inflammatory responses^28,29^. Further investigation into differentially expressed transcription factors in these cells showed uniquely higher expression of *SOX2* and *NRD1*, the latter previously associated with tumor-induced fibroblast activation^30^. Airway fibroblast progenitors expressed higher levels of *SERPINF1, FBN1, IGFBP5.* Based on the elevated expression of these genes, airway fibroblasts appear to emerge around GW15 and uniquely expressed the transcription factor *HOXA10* (**Fig. 2D**). Airway SMC differentially expressed higher levels of *TAGLN, ACTA2, MYH11*, while vascular SMC expressed *IGFBP7, MEF2C, EGFL6*, both similarly reported in other studies^9^. Transcription factors differentially expressed in airway and vascular SMC included the zinc finger E-box binding homeobox 1 (*ZEB1),* a core EMT transcription factor^11^, and the ETS variant transcription factor (*ETV1)*, which regulates cell growth, proliferation and differentiation, respectively. Chondrocytes differentially expressed many of the collagen genes including *COL2A1, COL11A1, COL9A*1 and the transcription factor CAMP Responsive Element Binding Protein 3 Like 2 (*CREB3L2* known regulate chondrocyte development. An unknown small cluster of stromal cells with distinct DEG expressed high levels of *AHSP, ALAS,* and *BPGN*, genes correlated with erythroid cells. They also expressed abundant levels of TMEM158, a gene enriched in cells undergoing epithelial-to-mesenchymal transition through STAT3 signalling^31^. The role of these unknown cells in fetal lung remains to be determined.

We performed spatial transcriptomics using 10X’ Visium technology and confirmed the localization of airway fibroblast progenitors, lipofibroblast precursors, and airway SMC in GW15 and 18 fetal lung tissues (**Extended Data** Fig. 2). Early fibroblasts, lipofibroblasts, lipofibroblast precursors, and airway SMC are broadly distributed throughout the GW15 and GW18 fetal lungs. In contrast, airway fibroblast progenitors are primarily concentrated around the large airways, a region where lipofibroblast precursors are distinctly absent. This spatial distribution suggests that early fibroblasts, lipofibroblasts, lipofibroblast precursors, and airway SMC contribute broadly to lung development, whereas airway fibroblast progenitors contribute specifically to airway development.

To infer developmental lineages of the fibroblast clusters, we used LatentVelo to estimate RNA velocities and visualized these velocities on the UMAP embedding and with PAGA (**Fig. 2E, F**). Inferred trajectories suggested that cycling fibroblasts form early fibroblasts and uncommitted cells that then contributed to airway fibroblast progenitors and lipofibroblasts. We further validated these trajectories with Slingshot, using cycling fibroblasts as the “root cells”, or cell origins and found a gradual change in gene expression as the cells along a differentiation trajectory to lipofibroblasts and airway fibroblast progenitors (**Fig. 2G, H**). To determine whether these trajectories changed over time, we grouped the datasets based on gestational time (Early: GW10-13, Mid: GW14-16 and Late: GW17-19) but found no changes in these trajectories when subsetting the analyses by GW (data not shown).

### A dynamic developing fetal lung epithelium

Analysis of the top DEG coupled with high expression of canonical epithelial genes associated with specific epithelial cell types revealed 19 subtypes (**Fig. 3A, B** and **Supplementary Table 2**). Many of the epithelial cell types were found in all lung tissues except for ciliated cell precursors which emerged around GW13 onwards. Mature ciliated cells and *SCGB3A2+FOXJ1+* cells emerged around GW14 whereas basal cells and SMG secretory cells emerged around GW15, but these latter cell types represented only a very small fraction (∼0.2-1% and ∼1-5%) of the total epithelial cell population (**Supplementary Table 3, Extended Data Fig. 3A**). Pulmonary neuroendocrine cells (PNEC) and *SOX2^high^CFTR+* cells were found in higher abundance in early fetal lungs (up to GW13) and then decreased in proportion in later gestational tissues.

**Fig 3:**
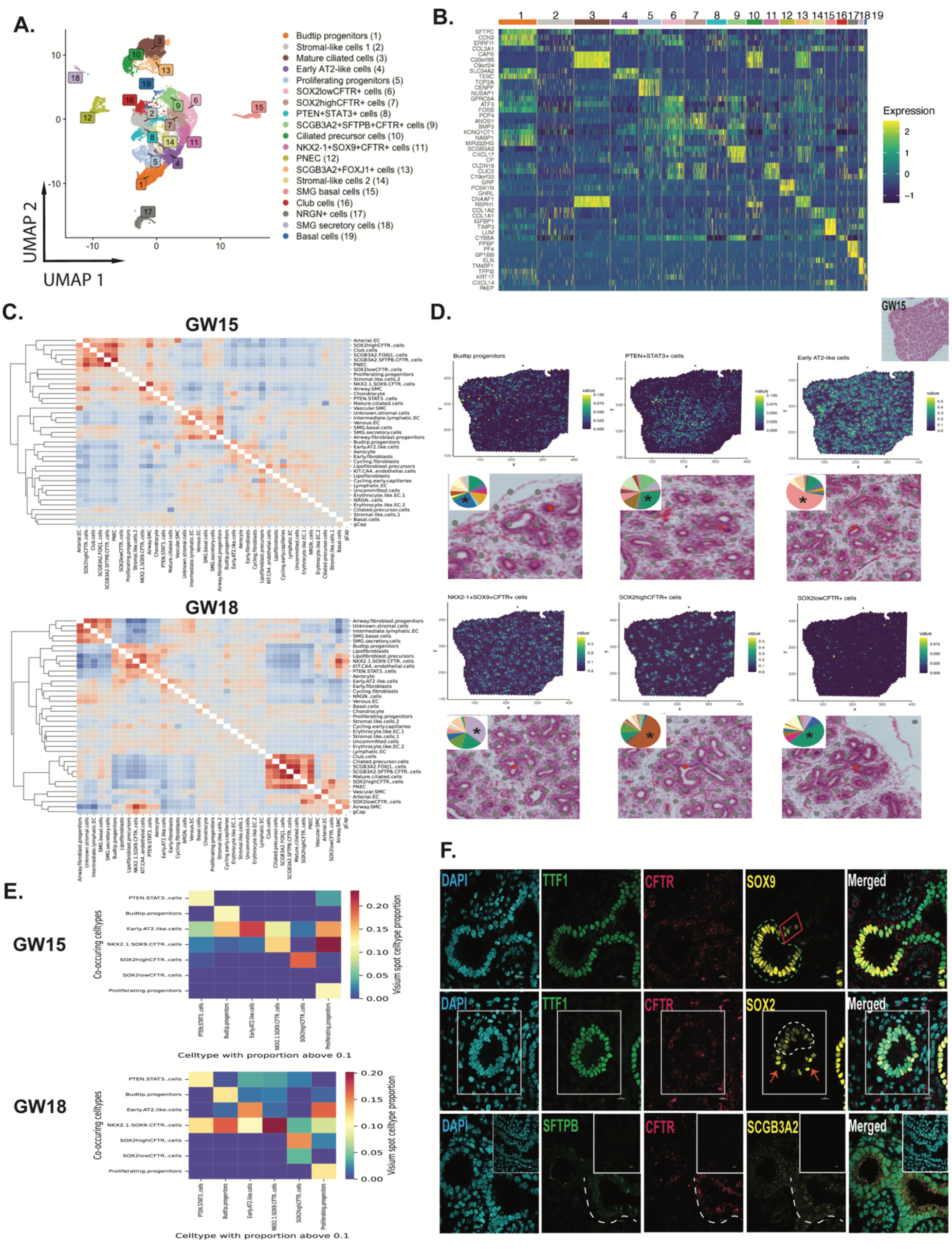
Heterogenous and novel cell types identified in the developing fetal lung epithelium. **A:** UMAP visualization of the fetal epithelial subtypes. **B:** Gene expression heatmap of the top 3 differentially expressed genes in each cluster. **C:** Correlation between RCTD weights for Visium spots. Showing the colocalization of specific celltypes within the same Visium spots. Hierarchical clustering shows groups of frequently colocalized cells. **D:** Spatial transcriptomic analysis (10X Visium) shows areas of localization of budtip progenitors, *PTEN+STAT3+* cells, early AT2-like cells, *NKX2-1+SOX9+CFTR+* cells, *SOX^low^CFTR+* cells, *SOX2^high^CFTR+* cells in GW15 fetal lung tissue. Location of Visium spots with highest RCTD weights (green-yellow hue) for the major immune cell types. High magnification of H&E staining shows a specific region where there is an abundant number of the specific cell subtype (red dot). Pie chart (insert) shows the proportion of the cells in the specific region (red dot). **E:** Given Visium spots with RCTD proportion for a specific cell type above 0.1, we plot the average proportion of spatially close epithelial cell types. **F:** Immunofluorescent staining shows the localization of SOX2, SOX9, CFTR, SCGB3A2, and SFTPB positive cells in the fetal lungs (GW15 tissue). Inset is the negative (no primary, secondary antibodies only) controls. DAPI marks all nuclei. Dashed lines and arrows demarcate areas of SOX2 low and high expression, respectively.

Leveraging the expression of *SOX2* and *SOX9* to determine proximal and distal airway cells respectively^32^, we identified subclusters that expressed relatively higher levels of *SOX2* and include PNEC, mature ciliated cells and their precursors, club cells, SC*GB3A2+FOXJ1+,* a CFTR- expressing cell type that co-expressed *SCGB3A2* and *SFTPB* (*SCGB3A2+SFTPB+CFTR+*^33^), and *SOX2^high^CFTR+* cell populations (**Extended Data Fig. 3B**). On the contrary, *SOX9^high^*cells included early alveolar type 2 (AT2)-like cells, budtip progenitors, *NKX2-1+SOX9+CFTR+* cells and both the stromal-like cell populations. Cells that expressed relatively lower levels of both *SOX2* and *SOX9* included submucosal gland (SMG) basal cells, SMG secretory cells, and a calmodulin-binding protein neurogranin *(NRGN)*-expressing cells. Interestingly, while *SOX2* and *SOX9* co-expressing cells have previously been identified in double positive distal tip lung progenitors^34^, the expression of *SOX2* in the budtip progenitors in our dataset is relatively low compared to *SOX9*. Instead, cells that appeared to express both *SOX2* and *SOX9* at relatively equal levels included *SOX2^low^CFTR+* cells, *PTEN+STAT3+* cells, basal cells, and the proliferating progenitors.

Proliferating progenitors were identified by the elevated expression of genes associated with cellular proliferation *MKI67,* and cell cycle genes encoding the microtubule-binding protein *CENPF, NUSAP1,* and *CDK1* (**Fig. 3B**). Proliferating progenitors expressed abundant levels of *BRCA1* and *FOXM1,* and high levels of *TEAD4*, a downstream effector of the Hippo pathway involved in lung development^35^. Other than abundant expression of genes associated with cell proliferation (*TOP2A* and *CENPF*), these cells are relatively uncommitted.

An unknown epithelial cell cluster was found that expressed high levels of mTOR-related *STAT3* and *PTEN* (*PTEN+STAT3+*); no other cell-type-specific genes were differentially expressed in these cells. In the developing mouse lungs, mTOR signaling is required for *SOX2* acquisition and conversion of *SOX9*+ distal progenitors to *SOX2*+ proximal cells^36^. It is unclear if these *PTEN+STAT3+* cells are equivalent to transitional cells of distal to proximal progenitors, however these cells express DEG associated with cell cycle progression such as nuclear factor kappa B (*NFKB1)*, NFKB subunit (*REL)* and IFN regulatory factor 1 *(IRF1)* (**Extended Data Fig. 3C**). Selecting Visium dots (spatial area) with high proportion of *PTEN+STAT3+* cells show co-occurrence (or close proximity) of these cells with *NKX2-1+SOX9+CFTR+* cells and AT2-like cells in GW15 tissues, and only with *NKX2-1+SOX9+CFTR+* in GW18 tissues, suggesting these cells may reside near the developing airway bud (**Fig. 3E**). *PTEN+STAT3+* cells also express abundant levels of thrombospondin 1 (*THBS1,* **Supplementary Table 2**), an extracellular glycoprotein involved in cell-matrix and cell-cell interactions. Immunostaining for THBS positivity showed abundant and uniform expression in the developing bud especially in the early (GW12) lung (**Extended Data Fig. 3D**). In later tissues (GW15 and GW18) expression of THBS1 were found on the apical membrane of some epithelial cells in the developing airways and along the basement membrane of the larger airways.

A novel finding was the identification of four epithelial cell types that expressed relatively high levels of *CFTR* (**Extended Data Fig. 3B**). These included *NKX2-1+SOX9+CFTR+*, SOX2^high^*CFTR+*, SOX2^low^*CFTR+* and *SCGB3A2+SFTPB+CFTR+* cell populations. The *SOX2^low^CFTR+* shared many common DEG with *SOX2^high^CFTR+* cells, however relative to *SOX2^high^CFTR+* cells*, SOX2^low^CFTR+* cells differentially expressed *GPRC5A* (a G protein-coupled receptor class C group 5 member A), involved in retinoic acid signaling, a pathway instrumental in lung development^37^ (**Fig. 3B** and **Supplementary Table 2**). Spatial transcriptomics showed an abundance of the Early AT2-like and *SOX2^high^CFTR+* in the developing airways at GW15 (pie chart inset shows the proportion of cell types in the spatial region demarcated in the red dot of the H&E stain which contained the predominant respective cell type marked with an asterisk, **Fig. 3D**). On the contrary, *SOX2^low^CFTR+* cells were rare and found in the developing stalks of the airways. In later tissues (GW18), *NKX2-1+SOX9+CFTR+* cells appeared to dominate all of the developing airways, likely due to the large expansion in the number of airways formed at this point (**Extended Data Fig. 4A**). It is important to note that within a spatial area (demarcated in the Visium as dots) there is a heterogenous proportion of cell types that reside in that “focal” area. Using immunofluorescence staining, we determined that the TTF1+ or (NKX2-1) SOX9+CFTR+ cells were found mainly in the developing lung bud region (green hatched area), and with SOX9^low^ cells were found in the developing stalk region (red boxed region) (**Fig. 3E**). On the contrary, TTF1+SOX2+CFTR+ cells were found in the fetal airways with distinct clusters of SOX2^high^ (orange arrow) and SOX2^low^ (white hatched area) cells. The role of CFTR in the developing lung is unclear. However, previous studies in porcine fetal lungs suggest a role of CFTR in branching morphogenesis^38^. Transcription factors (**Extended Data Fig. 3C**) highly enriched in *SOX2^high^CFTR+* cells included *RBPJ,* the transcriptional effector of NOTCH signalling^39^, and *FOXA2,* which mediates branching morphogenesis and differentiation^40^. Interestingly, these cells also express high levels of *CD47* (**Supplementary Table 2**), a cell surface molecule used to enrich for multipotent human induced pluripotent stem cells (hiPSC)-derived lung progenitors^41^. *NKX2-1+SOX9+CFTR+* cells expressed high levels of *CPM*, *SFTPA*, *FGFR2,* and *MUC1* and the transcription factor *SMAD6*. SMAD signalling is important in alveolar stem cell development^42^, which suggests *NKX2-1+SOX9+CFTR+* may be an ancestral source of alveolar cells, as has previously been suggested^43^. Indeed, *CPM* has previously been identified as a cell surface molecule used to enrich for fetal lung epithelial progenitors generated from hiPSC that can generate alveolospheres^44^.

The *SCGB3A2+SFTPB+CFTR+* cells were found scattered throughout the airways (white dashed region, **Fig. 3F**). The transcription factors enriched in *SCGB3A2+SFTPB+CFTR+* cells included *CREB3L2* and *RUNX2* (**Extended Data Fig. 3C**), the former previously shown to regulate secretory cells^45^ and the latter previously shown to be a regulator of goblet cell differentiation^46^. This would suggest that the *SCGB3A2+SFTPB+CFTR+* cells shared a lineage relationship with secretory cells^47^.

Unlike in the adult lungs^3,48^, fetal secretory, ciliated, and SMG cells expressed relatively low levels of *CFTR.* The rare ionocytes which express the highest levels of *CFTR* in postnatal lung tissues^48^ were not detected in any of the fetal epithelia examined here. While this does not preclude the development of these cells later in fetal lung development, the canonical genes (coexpression of *FOXI1* and *CFTR*) used to identify ionocytes could not be used to identify any distinct ionocyte-like populations. Fetal club cells expressed high levels of the canonical genes *SCGB1A1, SCGB3A2,* and *CYB5A* (**Supplementary Table 2**) typically found in mature club cells. On the contrary, *SCGB3A2+FOXJ1+* cells expressed abundant genes associated with ciliogenesis (*RSPH1, DNAH5*) cells and would suggest these may be transitory cells originating from a secretory precursor.

Ciliated cell precursors and mature ciliated cells shared expression of several key genes including *DNAH5,* a protein-coding gene for microtubule assembly, *FOXJ1*, a key regulator of cilia gene expression, and *RSPH1*, a gene encoding a protein that localizes cilia (Cluster 10, **Fig. 3B).** Both cell types also expressed high levels of the transcription factor *TP73* (**Extended Data Fig. 3C**), an activator of ciliary marker *FOXJ1*^49^. Ciliated cell precursors however expressed high levels of *FOXN4* (**Supplementary Table 2**), a transcription factor required for expression of motile cilia genes and observed in cells undergoing ciliated cell differentiation^50^. Mature ciliated cells, on the other hand, expressed prominent levels of the *TUBA1A* required for ciliogenesis^51^. Interestingly, we found a unique cell cluster that co-expressed *SCGB3A2* and *FOXJ1* (*SCGB3A2+FOXJ1+*) which were found dispersed in the airways (**Extended Data Fig. 3E**). The relationship between these cells with ciliated cells remains to be experimentally determined however assessment of the proximity scores of the cells from spatial transcriptomics suggest *SCGB3A2+FOXJ1+* cells are closely with associated with proximal cells including *SCGB3A2+SFTPB+CFTR+* and SOX2+ cells (**Fig. 3C**), suggests a lineage relationship between these cells. Below, we infer this relationship with RNA velocity.

Airway submucosal gland (SMG) basal cells expressed high levels of *IGFBP, TIMP3, LGALS1, LGALS3, S100A4,* and *S100A6,* all previously associated with SMG basal cells^9^. Similarly, SMG secretory cells expressed high levels of *UPK3B, RARRES2, C3,* and *ALDH1A2*. Both SMG basal and SMG secretory cells shared enriched expression of the *HOX* family transcription factors *HOXB5, HOXB2, HOXB4* and *HOXA5* (**Extended Data Fig. 3C**), all previously shown to regulate proximal-distal patterning and epithelial differentiation in the developing mouse airways^52^.

Early AT2-like cells and budtip progenitors both expressed genes related to surfactant protein production such as *SFTPC, SFTPA1,* and *ABCA3.* Budtip progenitors also expressed high levels of *CA2, TESC, ETV5* as previously identified^53^, in addition to *CCN1, CCN2*, *ITGA2*, genes associated with cellular proliferation *(EGR1*, *FOSB*), anti-apoptotic gene (*MCL1),* and *NFKBIA*. Budtip progenitors were found to be enriched with transcription factors *NFKB1* and *SOX9* (**Extended Data Fig. 3C**), both involved in distal epithelia differentiation^54^. Early AT2-like cells on the other hand, expressed high levels of *MFSD2A*, *CPM*, and *IGFBP7*. Transcription factors enriched in early AT2-like cells included *ETV1* and *HNF1B*, as previously described^55^ as well as *SOX9* and *FOXP2.* While ETV5 was previously shown to regulate human distal lung identity^11^ and mouse adult lung AT2 identity^56^, here we identify the homologous ETV1 as abundantly expressed in human fetal AT2-like cells. Proximity score analysis showed budtip progenitors and AT2-like cells were closely associated, suggesting these cells may also share regionally the same space (**Fig. 3C** and **3E**).

Other epithelial cell populations that were less defined and make up a smaller proportion of the total epithelium include the *NGRN+* cells that expressed abundant levels of *CCL5, TGFB1, GNG11,* which would suggest these cells may play a role in immunity and inflammation. Their precise role in development has yet to be determined. Two stromal-like epithelial cell populations were also found in all fetal lung tissues examined. These cells were annotated based on the DEG of stromal associated genes including *COL3A1* which is expressed in squamous cell types^57^. Stromal-like cells 2 also expressed the mesenchyme homeobox 2 (*MEOX2*)^58^ which regulates TGFbeta signalling pathway and epithelial-mesenchymal cell transition. These cells also express abundant *WNT2*, previously shown to orchestrate early lung morphogenesis^59^. Future studies will need to define the functional role of the stromal-like cells in the developing lung.

Similar to previous studies^9,10^, PNEC emerged early in the developing lungs but the proportion of PNEC appeared to decrease to relatively low proportions by the end of the pseudoglandular stage ∼GW16 (**Extended Data Fig. 3A**). PNEC expressed high levels of the canonical genes including *CHGB, CALCA,* and *ASCL1* (**Fig. 3B**) and the transcription factors *NEUROD1* and *ASCL1* were most abundantly expressed in these cells (**Extended Data Fig. 3C**), confirming their cell identity as previously described^9,10,60^. These cells were found in the airways (**Extended Data Fig. 4B**). Importantly, two PNEC subpopulations separated by differential expression of *ASCL1* and *CALCA* respectively was observed in our dataset (**Extended Data Fig. 4C**) and supports previous findings^9^.

In comparing the epithelial compartment to other recently published datasets of similar gestational ages, the ciliated cell precursors were not captured in the dataset by Peng et al.^9^, and Sountoudilis et al.^10^, (**Extended Data Fig. 4D** and 4E, respectively). Moreover, SMG secretory and both SMG and airway basal cell populations were not found in the Sountoudilis et al. dataset. This may reflect differences in clustering resolutions or cluster annotations.

Using LatentVelo, we predicted the lineage relationships of the epithelial cell types (**Fig. 4A**). We also used a separate pseudotime method to arrive at these same results for validation (**Extended Data Fig. 5A**). Analyzing LatentVelo velocities with *CellRank*^61^ reveals 3 terminal states: Mature ciliated cells, PNEC, and budtip cells **(Fig. 4B**, and **Extended Data Fig. 5B** and C). These identified terminal budtip cells are all at later GWs (GW 16-18). Using slingshot, we identified a bifurcation in the early budtip cells towards either the late budtip cells, or Early AT2- like cells and *NKX2-1+SOX9+CFTR+* cells (**Extended Data Fig. 5D**).

**Fig 4.**
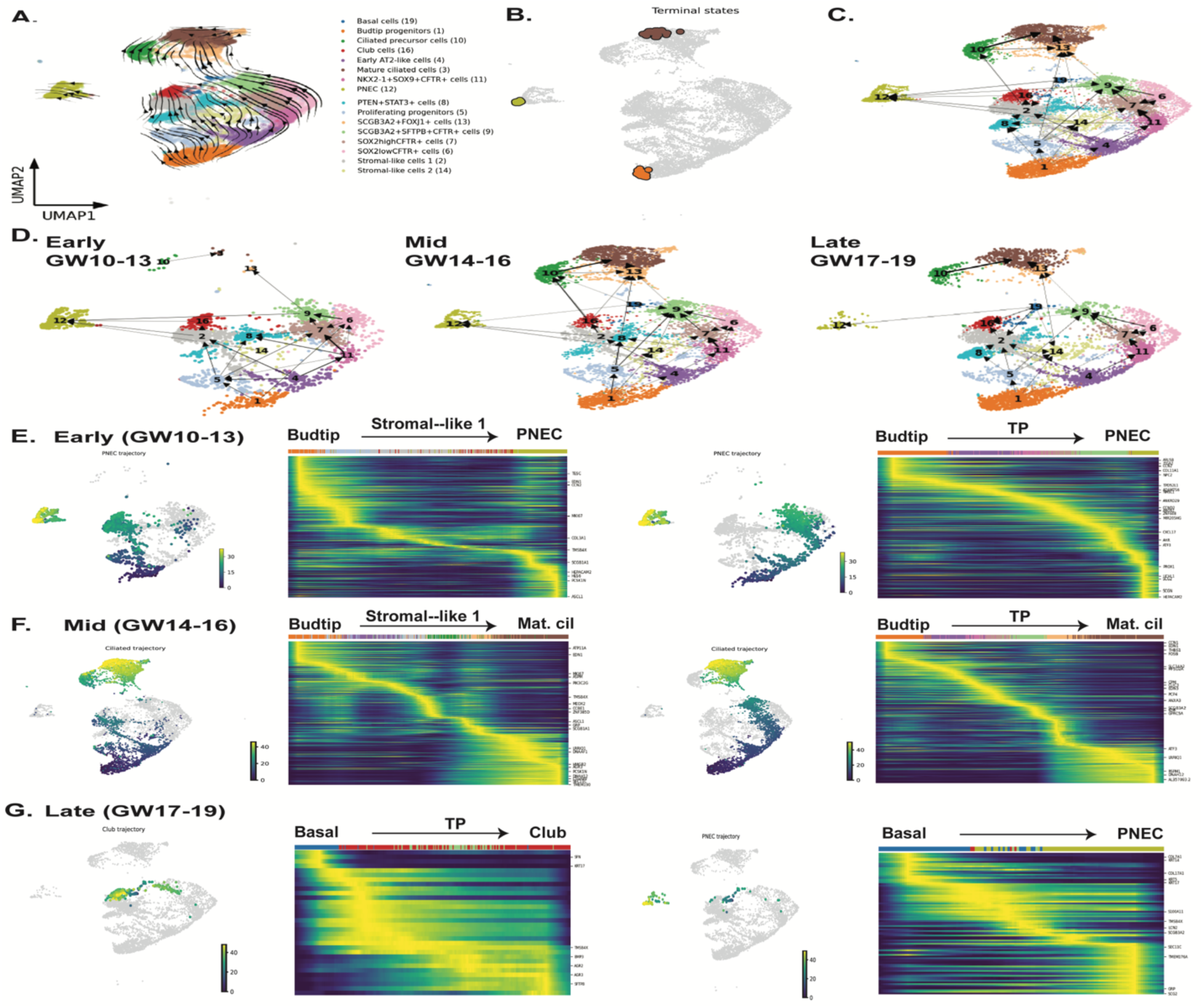
RNA velocity analysis reveals dynamic epithelial cell trajectories varying across gestational ages. **A:** LatentVelo analysis revealing inferred cell state trajectories projected onto the UMAP. **B:** Terminal states identified by using velocities with CellRank,shown on the UMAP. **C:** Partition-based graph abstraction (PAGA) analysis reveals multiple connections with the fetal epithelia. **D:** LatentVelo PAGA is subset into Early (GW10-13), Mid (GW14-16), and Late (GW17-19). The weakening of PNEC transitions were observed, and the emergence of transitions to ciliated cells with increasing GW. **E:** Based on the PAGA, the early trajectories to PNEC from Stromal-like cells 1 and *SCGB3A2+SFTPB+CFTR+* cells are analyzed with Slingshot. Trajectory heatmaps show the progression of cell types (colored bar) and significantly varying genes along the trajectory. **F:** Mid timepoint Slingshot trajectories highlighting the emergence of mature ciliated cells from Stromal-like cells 1 and *SCGB3A2+SFTPB+CFTR+* cells are shown. **G:** Based on PAGA, basal cells contribute to late club and PNEC cells. Slingshot trajectories show the development of PNEC cells from basal cells, and the development of Club cells from basal and *SCGB3A2+SFTPB+CFTR+* cells.

Based on PAGA using LatentVelo velocities, we observed trajectories that suggested a high degree of cellular plasticity of the proximal epithelial compartment as several trajectories would converge on the same cell types, in particular the identified terminal PNEC and mature ciliated states (**Fig. 4C**). We hypothesized that epithelial lineage development may be temporally regulated, and therefore to unmask these dynamic events, we needed to dissect the analysis based on developmental vignettes. Based on the emergence of several canonical cell types, such as mature ciliated cells and the significant decline of PNEC cells at ∼GW14 (**Extended Data Fig. 3A**), we decided to group the tissues into 3 vignettes: Early (GW10-13), Mid (GW14-16), and Late (GW17-19). The late vignette (GW17-19) marks the early canalicular stage of lung development, a period in which differentiation and emergence of specific cell types emerge. A few noteworthy trajectories were observed. First, in early gestational tissues, there were multiple sources of PNEC contributions including 1) a trajectory that goes through stromal-like 1 cells to PNEC; and 2) a trajectory from budtip progenitors through to *SOX2^high^CFTR* to *SCGB3A2+SFTPB+CFTR+* and then PNEC^33^. Changes in the DEG showed the gradual cell fate changes as the cells eventually gained expression of genes related to PNEC differentiation including *ASCL1* and *GHRL* (**Fig. 4E**). However, all of the PNEC trajectories weakened in the late stages. Several trajectories seemed to converge onto *SCGB3A2+SFTPB+CFTR+* in “mid” gestational tissues (**Fig. 4D**). These include cell sources from *SOX2^low^CFTR+* cells, *SOX2^high^CFTR+,* stromal-like cell 2, proliferating progenitors and basal cells. A strong connection (thick arrow line) between *NKX2- 1+SOX9+CFTR+* and *SOX2^high^CFTR+* cells and the latter with *SCGB3A2+SFTPB+CFTR+* were observed in all timepoints, suggesting these cells are developmentally related. While a connection was observed between *NKX2-1+SOX9+CFTR+* and *SOX2^low^CFTR+* cells, this weakened in late tissues. Similarly, a connection between budtip progenitors and *SOX2^low^CFTR+* cells were also observed in all timepoints. Interestingly, the connection between budtip to AT2-like cells appeared to reflect only a subpopulation of budtip cells that emerged later in development (mid stage). During the mid (GW14-16) timepoints, the emergence and trajectories for mature ciliated cells were observed, stemming from budtip progenitors through stromal-like 1, club, and ciliated precursor cells or through *SCGB3A2+SFTPB+CFTR+* cells and *SCGB3A2+FOXJ1+* cells (**Fig. 4F**). Fetal club cells appeared to contribute to the ciliated cell precursor (**Fig. 4D**) supporting the notion that Club cells may have greater developmental potential as has been observed in mouse lung injury models^62^. Interestingly, there were no connections between fetal basal cells and club cells before GW17, which may be explained by the low numbers of basal cell captured for our analysis. In late tissues, trajectories originating from basal cells contribute to PNEC and Club cell lineages (**Fig. 4G**), suggesting the basal cells at this stage have acquired developmental plasticity similar to that observed in adult basal cells^63^.

### The plasticity of SCGB3A2+SFTPB+CFTR+ cells in PNEC and ciliated cell lineages

Similar to Conchola et al^33^, we also identified a *SCGB3A2+SFTPB+CFTR+* population with a developmental contribution to PNEC and ciliated cells, but also Club cells in later gestational tissues (>GW16). By resolving temporal dynamics, we found a decline in PNEC contribution and concomitant increase in ciliated cell contribution by *SCGB3A2+SFTPB+CFTR+* cells at gestational week 14 (**Fig. 5A**).

**Fig 5.**
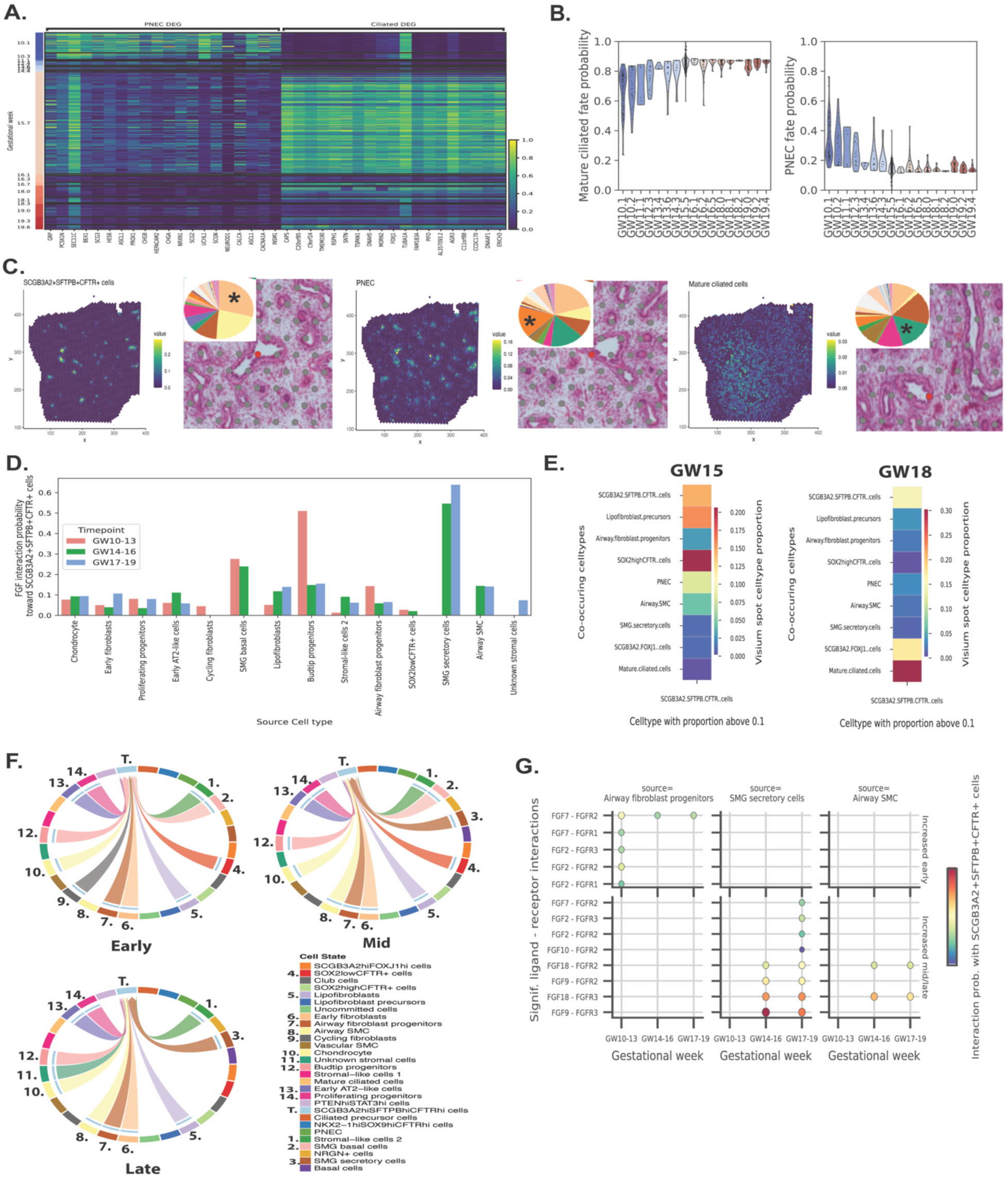
Regulation of TP cells differentiation and cell communication. **A:** Heatmap showing the gene-normalized expression of the top 20 differentially expressed genes for the PNEC and mature ciliated cells within the *SCGB3A2+SFTPB+CFTR+* cells versus Gestational Week. **B:** CellRank mature ciliated and PNEC fate probabilities for *SCGB3A2+SFTPB+CFTR+* cells versus Gestational Week. **C:** Spatial transcriptomic analysis (10X Visium) shows areas of localization of *SCGB3A2+SFTPB+CFTR+* cells, PNEC, mature ciliated cells in GW15 fetal lung tissue. Location of Visium spots with highest RCTD weights are demarcated in green-yellow hue. High magnification of H&E staining shows a specific region where there is an abundant number of the specific cell subtype (asterisk). Pie chart (insert) shows the proportion of the cells in the specific region (red dot). **D:** FGF signalling interaction probabilities towards *SCGB3A2+SFTPB+CFTR+* cells from specific cell types. **E:** Given Visium spots with RCTD proportion for *SCGB3A2+SFTPB+CFTR+* cells above 0.1, we plot the average proportion of spatially close cell types. **F:** FGF signalling interactions towards *SCGB3A2+SFTPB+CFTR+* cells subset to cell types in close proximity (Fig. 5E). Size of the incoming arrows indicate strength of signalling. **G**: Significant ligand-receptor (L-R) interactions towards *SCGB3A2+SFTPB+CFTR+* cells for the FGF pathway, subset to signalling cell types in proximity. Both color and point size indicate interaction probability. Top row shows L-R interactions significantly increased early. Bottom row shows interactions significantly increased mid/late. All significance tests use p < 0.01.

We used CellRank to estimate the PNEC or mature ciliated cell fate probability for the *SCGB3A2+SFTPB+CFTR+* cells, showing a decline in PNEC probability and increase in mature ciliated cell probability with GW (**Fig. 5B**). Spatial transcriptomics showed the proximity of both PNEC and ciliated cells with *SCGB3A2+SFTPB+CFTR+* as evident in the demarcated region (red dot) of the scatterpie plots of cell proportions (**Fig. 5C**). Within Visium spots, spatial proximity of PNEC and *SCGB3A2+SFTPB+CFTR+* cells were identified at week 15, while spatial proximity of mature ciliated cells, *SCGB3A2+FOXJ1+* cells, and *SCGB3A2+SFTPB+CFTR+* cells were found at week 18 **(Fig. 3C and 5E)**. Together, these results demonstrate a temporal regulation of PNEC and ciliated cell differentiation through *SCGB3A2+SFTPB+CFTR+* cells, supporting the dynamic plasticity of these progenitor cells in the developing lung.

### Temporal changes in cell signalling regulates SCGB3A2+SFTPB+CFTR+

Several new trajectories were observed in mid and late tissues which would suggest that cell fate acquisition is much more dynamic than previously understood. One explanation for these dynamic cell fate changes may reflect changes in the spatial microenvironment of the cell^64^ and the various signalling mechanisms (paracrine, autocrine, distant signalling)^65^. To determine the cellular interactions in regulating the change in gene expression and developmental potential of *SCGB3A2+SFTPB+CFTR+* cells, we performed *CellChat*^62^ analysis on our scRNA-seq dataset and identified unique signaling pathways targeting the *SCGB3A2+SFTPB+CFTR+* cells and observed changes in signalling when assessed through each vignettes: early (GW 10-13), mid (GW 14-16), and late (GW 18-19) (**Extended Data Fig. 6A**,B). Signaling pathways that specifically targeted *SCGB3A2+SFTPB+CFTR+* cells relative to other epithelial cells, included fibroblast growth factor (FGF) and WNT signaling. We also observed high levels of NOTCH signaling in early-mid tissues that then decreased in mid-late tissues (**Extended Data Fig. 6A**, B).

We then focused our investigation on the FGF signaling pathway as it was observed to have significantly increased (red arrow, **Extended Data Fig. 6A**) over time and known to play a role in branching morphogenesis during lung development^66^. Moreover certain FGF recombinant proteins are used in many directed differentiation protocols to generate lung epithelia from human iPSC^13,15^. We specifically focused on cell-cell interactions through FGF signaling with *SCGB3A2+SFTPB+CFTR+* cells **(Fig. 5D**). Informed by Visium, we identified cells that were spatially in “close proximity” to *SCGB3A2+SFTPB+CFTR+* cells which included lipofibroblast precursors, *SOX2^high^CFTR+* cells, PNEC and airway SMC in GW15 lungs; and *SCGB3A2+FOXJ1+* and mature ciliated cells in GW18 lungs (**Fig. 5E**). Further analysis of the potential signaling partners involved in FGF signaling in *SCGB3A2+SFTPB+CFTR+* cells (receiver cells of the signal) across time, showed multiple contributions of sender cells that are near and cells that are not spatially close to *SCGB3A2+SFTPB+CFTR+* cells (**Fig. 5F**). Interestingly, there were temporal changes in FGF sending cells, but there were also persistent signals from the same senders (ie. Airway fibroblast progenitors) that remained throughout the time points. Importantly, we did not include PNEC, *SCGB3A2+FOXJ1+*, or mature ciliated cells (which are also spatially close to *SCGB3A2+SFTPB+CFTR+* cells), as we aimed to find the signals that may contribute to the dynamic contribution of *SCGB3A2+SFTPB+CFTR+* cells to PNEC/ciliated cells.

We then estimated the interaction probability between specific ligand-receptor (L-R) interactions involved in FGF signaling across GWs between *SCGB3A2+SFTPB+CFTR+* cells and FGF signal senders, specifically airway fibroblast progenitors, SMG secretory and airway SMC as they were determined to be cells in close proximity with SCGB3A2+SFTPB+CFTR+ cells, based on Visium spatial transcriptomics (**Fig. 5E, 5G**). We found significantly increased signaling between L-R interactions involving the ligands FGF2 and FGF7 expressed in airway fibroblast progenitors during GW10-13, and we found significantly increased signaling for L-R interactions involving the ligands FGF9, FGF10, and FGF18 expressed in SMG secretory and airway SMC during GW14-16 or GW17-19. Gene expression of the specific FGF ligands and receptors showed elevated expression of FGF receptor 2 (*FGFR2*) and 3 (*FGFR3*) in *SCGB3A2+SFTPB+CFTR+* cells **(Extended Data Fig. 6C).** We confirmed the high expression of FGFR2 and FGFR3 using Visium, in regions with high proportions of *SCGB3A2+SFTPB+CFTR+* cells at GW 15 and 18 (black box, **Extended Data Fig. 6D).** These results suggest greater contribution of FGF2 and FGF7 signaling by airway fibroblast progenitors during early development, and FGF9, FGF10, and FGF18 signaling by SMG secretory and airway SMC later in development and may explain the temporal switch from PNEC to ciliated cells contributions.

We also assessed NOTCH signaling in *SCGB3A2+SFTPB+CFTR+* cells and found relatively strong NOTCH signaling in lung tissues during the mid (GW14-16) timepoint **(Extended Data Fig. 6B).** NOTCH receptors: *NOTCH2* and *NOTCH3* were expressed in multiple cell types including *SCGB3A2+SFTPB+CFTR+* cells and *SCGB3A2+FOXJ1+* cells, but not in PNEC **(Extended Data Fig. 7A**). Whereas NOTCH ligand *JAG1* was expressed in multiple cell types including vascular SMC and *SOX2^high^CFTR+* cells. High **e**xpression of *NOTCH2* and *NOTCH3* was found in spots with high proportion of *SCGB3A2+SFTPB+CFTR+* cells (black box, **Extended Data Fig. 7B**). Several sources of NOTCH signaling with *SCGB3A2+SFTPB+CFTR+* cells included vascular SMC, *SOX2^low^CFTR+* cells, and *SOX2^high^CFTR+* cells **(Extended Data Fig. 7C**). Interestingly, airway SMC was a source of NOTCH signaling during the “early” stage that was lost in mid/late stages. Increased *JAG1- NOTCH2* and *JAG1-NOTCH3* L-R interactions are also observed in “mid” stage tissues stemming from *SOX2^high^CFTR+* cells (**Extended Data Fig. 7D**). Overall, our data suggests NOTCH-mediated signaling to *SCGB3A2+SFTPB+CFTR+* during “mid” stage may involve direct interaction with *SOX2^high^CFTR+*.

WNT signaling in *SCGB3A2+SFTPB+CFTR+* cells were also assessed and found to be elevated in early (GW 10-13) and decreased in mid and late stages **(Extended Data Fig. 6B).** WNT receptors *FZD2, FZD3, FZD5, FZD6, FZD7, LRP5, LRP6* were found to be expressed in *SCGB3A2+SFTPB+CFTR+* cells with comparatively higher expression of *FZD2* and *FZD7* in the early stages **(Extended Data Fig. 8A**). Based on spatial expression from Visium, we identified *FZD2, LRP5, LRP6,* and *FZD7* expressed in areas of abundant *SCGB3A2+SFTPB+CFTR+* spots at GW15 whereas *LRP5, FZD3, LRP6,* and *FZD6* were expressed at GW18 **(Extended Data Fig. 8B**). Sources of WNT signalling was assessed in early, mid, late stages to *SCGB3A2+SFTPB+CFTR+* cells where *WNT2* was derived from multiple cell types including airway SMC (only in early stage), lipofibroblast precursors, and airway fibroblast progenitors (**Extended Data Fig. 8C, 8D**). Interestingly, signals from SMG secretory cells appeared to dominate in mid/late tissues (**Extended Data Fig. 8D**).

While all of these pathways have previously been described in mammalian lung development^67^, future studies will need to confirm their explicit role in specific cell lineage development and how the cell interprets the wide array of signals it receives to effect specific developmental functions.

### Human pluripotent stem cell (hPSC)-derived fetal lung cultures capture cellular heterogeneity and trajectories found in the native tissue

Directed differentiation protocols of hPSC aim to capture developmental milestones ensuring robust development of bona fide cell types in cell cultures. However, to do this, differentiation protocols must reflect developmental processes that are observed in the primary tissues. Many differentiation protocols, including for the lung, captures “fetal-like” states but few have benchmarked the developmental stage of these cells to fully understand the phenotype of the cells and potential limitations of the protocols. The human “fetal” stages of tissue development span nearly 9 months of gestation, during which a small batch of early embryonic cells must proliferate, differentiate and migrate to form a complex functional tissue. As the differentiations become more advanced, with the capability to generate hPSC-derived tissues-specific organoids, these models represent an exciting research tool to study fundamental mechanisms of development, especially when access to primary fetal tissues for research is limited. Here, we sought to determine the developmental stage of the cells we considered to be “fetal” lung cells and organoids generated from hPSC using our most recent lung differentiation protocol^13^ (**Figure 6A**). We performed scRNA-seq on the hPSC-derived fetal lung cells, considered to contain mostly undifferentiated lung epithelial cells, and fetal organoids, which were subjected to further differentiation. Using batch-balanced nearest neighbors, we integrate the hPSC-derived cells and the primary fetal lung epithelia. The integrated clustering showed significant overlap of hPSC-derived fetal lung cells (green cluster) and organoids (purple cluster) with epithelial cells in the primary tissue (**Fig. 6B, C**). A Pearson correlation coefficient between the hPSC derived cultures and the tissues across gestational timepoints showed a strong correlation (>0.75) to GW13 lung epithelial for hPSC-derived fetal lung cells (Stage 3), and (>0.75) to GW16 to 19 lung epithelia for hPSC-derived organoids (**Fig. 6D**) which represent the next stage of differentiation (Stage 4, **Fig. 6A**). To unbiasedly determine the epithelial cell subtype in the hPSC-derived cultures, we created cell type scores from the top 100 DEG for each cluster identified in the primary epithelial tissue (**Fig. 6E**). Interestingly, proliferating progenitors were the greatest scoring cell type in the hPSC-derived fetal lung cells. This is not surprising as the culture conditions was intended for fetal lung cell growth including retinoic acid, CHIR99021, and FGF7^53^. Other cell types with elevated scores included PNEC, basal, SMG basal and secretory, and stromal-like cells. On the contrary, hPSC-derived fetal lung organoids which represents the next stage following expansion of the fetal lung cells, contained high basal cell scores, along with club cells, PNEC, *SCGB3A2+SFTPB+CFTR+*, SMG basal and secretory cells, and stromal-like cells. The high basal cell score was not surprising as our organoid expansion media contained dual TGFbeta/SMAD signalling inhibitors DMH1 and A83-01 which promotes basal cell expansion^68^. To understand the extent to which these hPSC organoids represent fetal lung epithelial development, we positioned these cells along our previously inferred trajectories from the native tissue. We considered the previously inferred trajectory for the primary fetal epithelium from Budtip progenitors to *SCGB3A2+SFTPB+CFTR+* cells, through early AT2-like cells, *NKX2-1+SOX9+CFTR+* cells, and S*OX2^high/low^CFTR+* cells (**Fig. 6F**). Using the integrated nearest neighbors graph we clustered the integrated cells and only assessed the clusters with at least 100 cells along the trajectory (**Fig. 6G**). In particular, the hPSC-derived cells resided in clusters 1, 9, 7, and 5, with hPSC organoids in all 4 of these clusters and hPSC fetal lung cells only within cluster 1 **(Fig. 6H)**.

**Fig 6:**
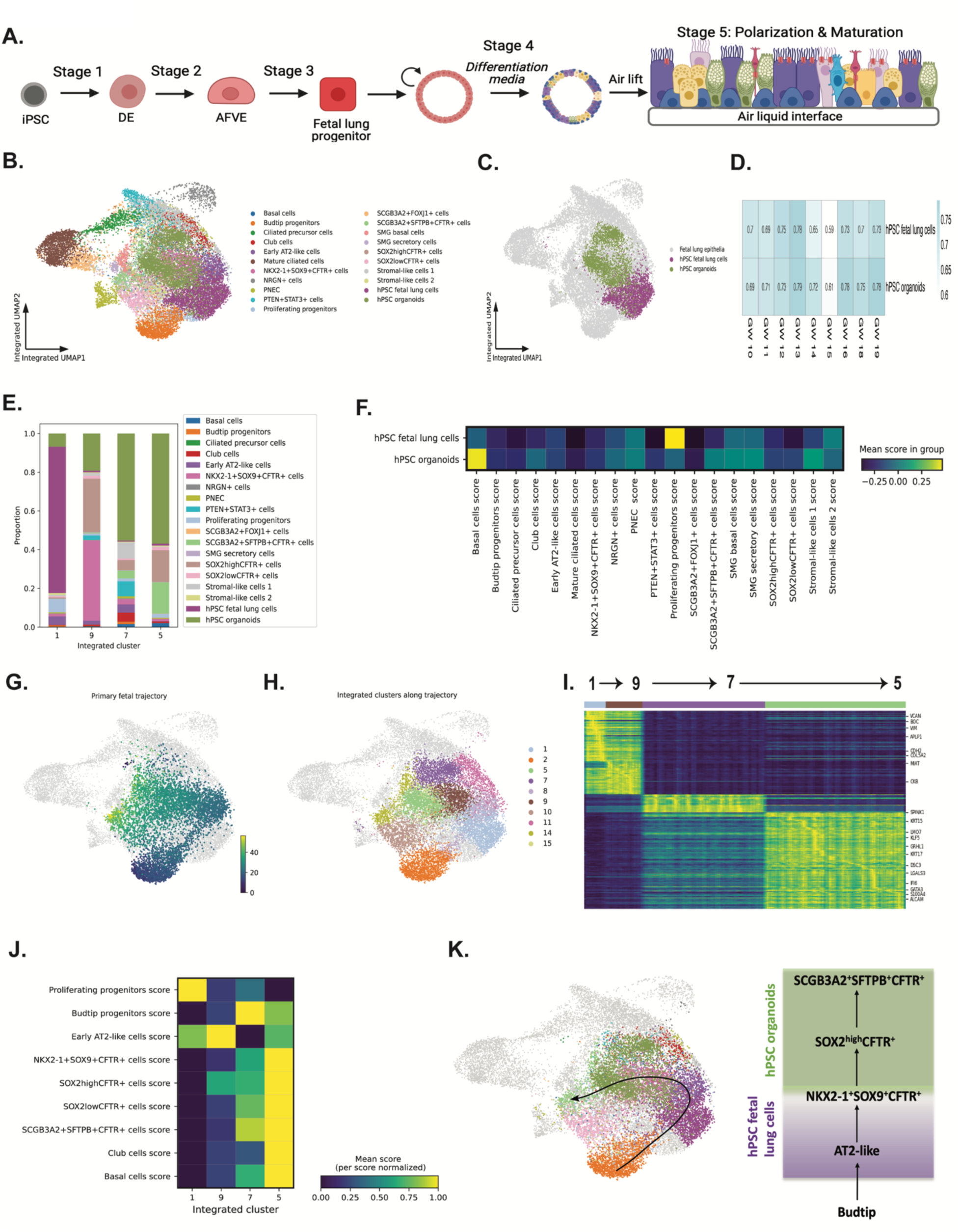
hPSC-derived lung models recapitulate trajectories towards basal and CFTR- expressing cells from the fetal lung epithelia. **A:** Stage-specific differentiation of hPSC towards mature proximal airway epithelia. **B:** Integrated UMAP projection of hPSC-derived fetal lung cells and organoid integrated with primary fetal lung epithelia. **C:** Integrated UMAP projection of hPSC-derived fetal lung cells and organoids cells. **D:** Pearson correlation between hPSC-derived fetal lung models and primary-derived fetal lung epithelia using the top 50 DEG for each gestational week. **E:** Proportion of primary and hPSC derived fetal models within clusters 1, 5, 7, and 9. **F:** Cell type scores based on the top 100 DEGs from the primary fetal epithelia reveal high proliferating progenitor scores in the hPSC fetal lung cells and high basal scores in the hPSC organoids. **G:** Previously derived primary fetal trajectory from Budtip progenitors to *SCGB3A2+SFTPB+CFTR+* cells through Early AT2-like, *NKX2-1+SOX9+CFTR+*, and *SOX2^high^CFTR+* cells on the integrated UMAP. Color shows increasing pseudotime. **H:** Integrated clustering of the primary fetal and hPSC derived cells focused on clusters 1, 5, 7 and 9 which contains hPSC organoids and primary fetal cells. **I:** Trajectory heat map showing the change in gene expression of hPSC organoids within these clusters, ordered by increasing average pseudotime of the primary fetal cells within each cluster. **J:** Celltype score for the hPSC organoids within the ordered clusters. **K:** Primary fetal and hPSC derived cells along the inferred trajetory. hPSC fetal lung cells resemble the early part of the trajectory, and hPSC organoids the later.

Using the average pseudotime of each of these clusters containing hPSC organoid cells, we plotted a heat map of significantly changing genes along the trajectory. A clear pattern of the change in gene expression emerges, and an analysis of the cell type scores **(Fig. 6I)** indicated that these organoids become increasingly differentiated along this trajectory with the acquisition of specific genes associated various differentiated epithelial cell types/states. In particular, proliferating progenitors and early AT2-like cell type scores were lost with increasing pseudotime and *SOX2^high/low^CFTR+*, *SCGB3A2+SFTPB+CFTR+*, Basal, and club cell type scores increased (**Fig. 6J**). Overall, our hPSC differentiated fetal cell models captures some of the epithelial cell types/states and trajectories observed in the fetal pseudoglandular/canalicular lung.

## Discussion

Reactivation of developmental mechanisms occur during disease pathogenesis^69^(such as pulmonary fibrosis) and tissue repair^70^. Therefore, understanding the plasticity of cells under normal development and the role of the local cellular signalling environment in dictating cell fate and function may provide key insights into congenital lung diseases, chronic disease pathogenesis and the cellular responses to therapies. Here, we created a comprehensive topographic human fetal lung transcriptomic atlas of the developing human fetal lung. Leveraging over 150,000 fetal cells captured, we identify novel cell types/states and inferred developmental trajectories revealing remarkable cellular plasticity within the epithelial compartment. With spatially resolved transcriptomics, we identify putative cell signalling interactions that dictate cell fate and behaviours within the developing lung. Finally, we show for the first time, the differentiation of hPSC progressing along similar developmental trajectories and giving rise to similar fetal cell types/states as observed in the human fetal lungs. This demonstrates the potential of the hPSC as experimentally tractable models in elucidating human-specific developmental mechanisms.

The pseudoglandular lung is marked by extensive branching morphogenesis, where the cells must undergo extensive proliferation, proximal-distal patterning, cell migration and differentiation into specialized cell types. A major novel finding in our analysis is the identification of several progenitor cells expressing high levels of CFTR. These cells include the *NKX2- 1+SOX9+CFTR+, SOX2^high^CFTR+, SOX2^low^CFTR+, and SCGB3A2+SFTPB+CFTR+* cells. The co-expression of CFTR in these progenitor cell types suggest a putative role for CFTR in branching morphogenesis and the formation of specific epithelial cell lineages. In the postnatal lung, mutations in the CFTR gene can cause cystic fibrosis (CF) disease, a disease that manifests in the airways and impairs lung function and ultimate destruction. The *CFTR* gene encodes for a chloride channel that regulates water and ion transport across the epithelium. The precise role of CFTR in the developing lung is unclear but the early expression of CFTR may suggest a role for this protein and the cells that expresses it in airway formation. Moreover, understanding the role of CFTR in airway cell development may provide important insight into the early manifestations of CF lung disease and the long-term impact on disease pathogenesis.

We also uncover a broader epithelial cell plasticity and with inference of lineage trajectories involving the *CFTR+* cells. Proceeding from *NKX2-1+SOX9+CFTR+*, we found a trajectory through *SOX2^high^CFTR+* and *SOX2^low^CFTR+* cells to *SCGB3A2+SFTPB+CFTR+* cells. We also found that the *SCGB3A2+SFTPB+CFTR+* cells have the developmental potential for club cells (a secretory cell type), and both PNEC and ciliated cells as has previously shown with *in-vitro* lineage tracing^33^. We were also able to resolve the role of time in the contribution of various epithelial cell subtypes, in which the contribution to cell states/types changed over the course of the 10 weeks of gestation. We demonstrate the multi-lineage plasticity of the CFTR- expressing population and the remarkably dynamic developmental potential of these cells. Interestingly, cellular heterogeneity and high developmental plasticity has previously been shown in endothelial lineage development^71^ and neutrophil development^72^. Moreover, epithelial plasticity has also been shown to drive the formation of the endoderm during gastrulation^73^. Therefore, it is conceivable that the epithelial cells of the developing lung retained some of the high developmental plasticity that are not observed in more developed organs at the same timepoint. Notably, this dynamic plasticity was not observed in the stromal compartment. In comparing to published datasets from He et al^9^ and Sountoulidis et al.^10^, both found trajectories involving *SCGB3A2+* cells or “proximal progenitor cells” to PNEC respectively, but neither identified the *SCGB3A2+SFTPB+CFTR+* cells specifically to be the source, nor have they identified the various CFTR+ cell subsets that appear to contribute to many sub-epithelial cell types. Unlike Conchela et al.^33^, we show that contribution to mature ciliated cells from *SCGB3A2+SFTPB+CFTR+* involves *SCGB3A2+FOXJ1+* cells, which may be an intermediary transient cell state. Overall, we highlight how specific epithelial cell lineages are formed in the developing lungs which may inform new directed differentiation protocols or cell regeneration strategies for lung diseases.

Developmental processes contributing to cell fate decisions are mediated by precise cell-cell signaling interactions within a cell’s microenvironment. While cells may share transcriptomic gene signatures, cell autonomous and/or non-cell autonomous signalling may dictate how these cells develop. Here we focused on signalling communications between epithelial-epithelial and epithelial-stromal cells however, it does not preclude the contributions of the immune, endothelial, and neural (Schwann) cells that are also co-developed and residing near the former two cell types. Interestingly, *CellChat* analysis informed from spatial Visium data showed temporal change in the signaling pathways targeting the *SCGB3A2+SFTPB+CFTR+* cells, which would suggest that both spatial and temporal effects can influence cell behaviour and potential cell fate decisions. In our study, we confirmed FGF, WNT and NOTCH signalling in fetal lung development. Previous studies have demonstrated all three pathways to play a role in lung branching morphogenesis and epithelial differentiation^74^. Mouse studies have suggested distinct signalling pathways driving epithelial specification and branching via Fgf9 and Fgf10, respectively^75^. Moreover, lateral inhibition of Notch signalling increase PNEC differentiation in Hes1-null mice^76^. Similarly, in our data set, we observed a dramatic decrease in PNEC in later gestational lung tissues as NOTCH signalling temporally increased. Future studies will need to determine the precise role of all these signaling pathways (in addition to other pathways) in regulating specific epithelial cell differentiation. Altogether our data highlights the intricate nature of cellular communications within a cell’s microenvironment that may contribute to the heterogenous and complex development of the fetal lung epithelium.

An important aspect of our study was the benchmarking of the hPSC-derived fetal lung cells in which the fetal epithelial models generated^13^ can capture several of the fetal-specific cell types/states and differentiation trajectories observed in the native tissue. Using single-cell RNA sequencing, we found that differentiated hPSC-derived fetal lung cells, which mostly represents proliferative, uncommitted cells, and hPSC-derived fetal organoids, which represent a later developmental stage, shared similar transcriptomic expression to fetal epithelial cells in the pseudoglandular stage (GW13 and GW16-19, respectively). Since not all cell types and trajectories were captured in the hPSC differentiation, future studies will need to leverage the predictions of the L-R interactions to modify current differentiation protocols that may help improve the generation of other cell types in-vitro. Nonetheless, our work supports the use of hPSC-derived fetal lung epithelial models as a surrogate to study certain aspects of lung development. Moreover, hPSC differentiations are experimentally tractable models and can be used to study broader developmental ranges and origins of diseases, especially since primary tissues may be limited due or restricted.

Our extensive analysis of the developing human fetal lungs does have a few limitations. First, while recent publications^9,10^ have sequenced earlier fetal lungs (as early as 5PCW), our study does not capture these earlier timepoint tissues which could have uncovered additional heterogeneities. Second, our study focused on collecting and analyzing freshly isolated lung tissues in which the tracheas were manually removed and whole lungs immediately processed for single-cell library prep with very minimal processing/transfer time. No cell enrichment for specific populations were performed for the purpose of capturing all cell types. However, this may have inadvertently limited our ability to detect rare cells. Moreover, due to the technical difficulties of accurately discerning “proximal” versus “distal” airways, our data could not resolve regional differences between proximal vs distal specific cell types, as has previously been shown^9^. Despite these technical differences, our data unbiasedly captured many of the cell types/states and developmental trajectories previously observed in recent datasets^9,10,33^. Finally, while our data aimed to prevent over-clustering by using *Clustree* to identify the optimal resolution for sub- clustering, it is possible that this method may not have resolved finer differences in cell states resulting in lower cell types/states identified. Future studies aimed at combining our dataset with proteomics and perturbing gene regulatory networks in functional genomics using hPSC models would provide a powerful tool to validate the developmental regulation of the human fetal lung.

Overall, our study identified novel cell types/states, trajectories and local interactomes that contributes to the dynamic changes in the early human fetal lung epithelium. Understanding human fetal lung cell diversity and their role in development will improve current differentiation protocols for hPSC to generate better organoid models/bona fide cell types for future therapies for congenital and chronic lung diseases.

## Extended Data Figures

**Extended Data Fig 1:**
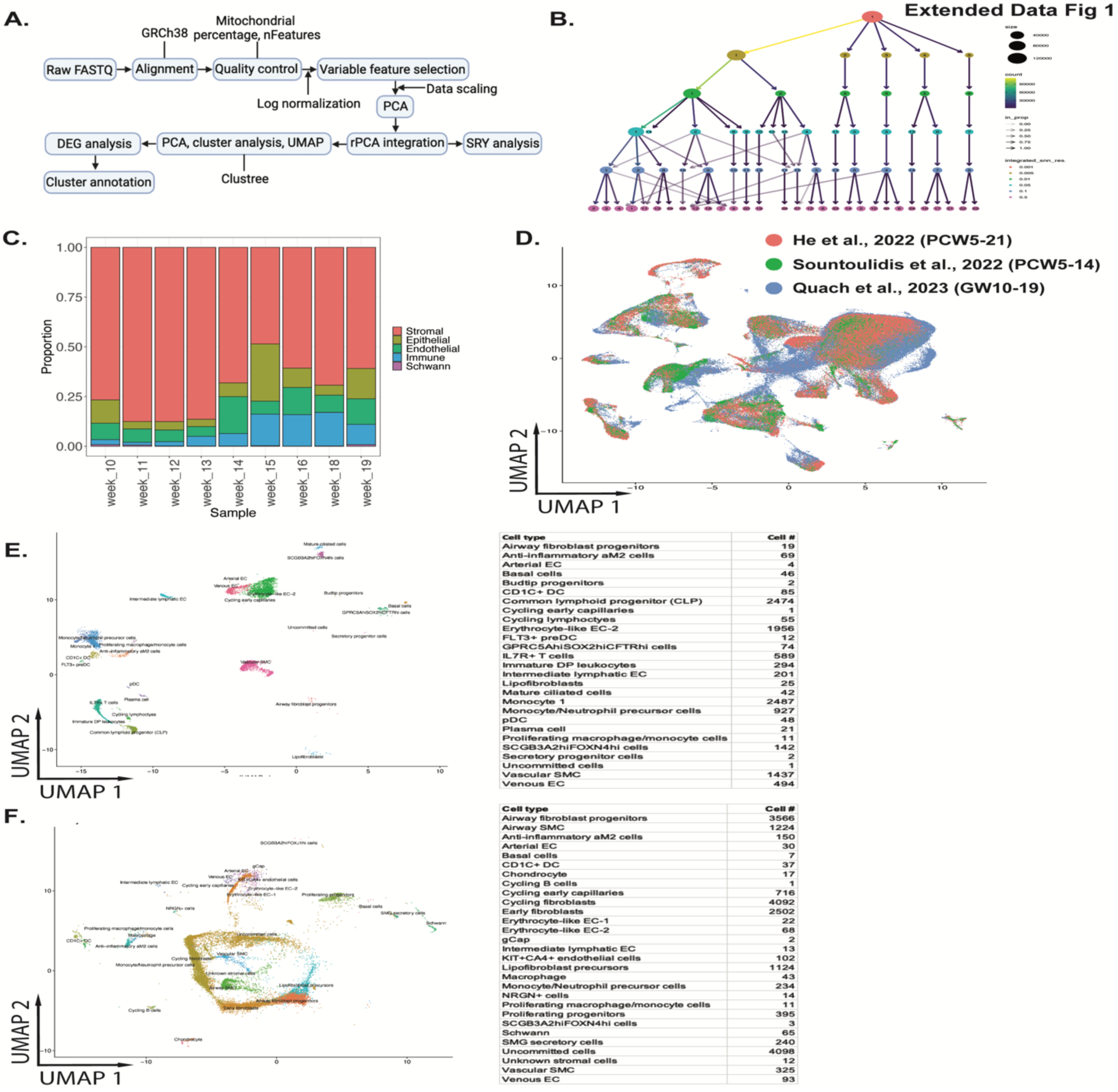
Analytical pipeline and characterization of the human fetal lung. **A:** Computational pipeline for single-cell RNA sequencing analysis. **B:** *Clustree* analysis to determine the number of cell clusters. We selected the resolution in which the number of cell clusters would not yield “overlapping” subpopulations. **C:** Proportion of the major cell types across GW. **D:** Integrated UMAP projections of all three datasets of human fetal lungs (He et al., 2022, Sountoulidis et al., 2022, and our dataset). **E:** *MapQuery* analysis using our dataset as the reference map to identify cell subtypes common in adult lung tissues. Adult lung datatset was obtained from Travaglini et al. 2020. **F:** *MapQuery* analysis using our dataset as the reference map to identify cell subtypes common in mouse embryonic lung tissues. Mouse lung datatset was obtained from Negretti et al. 2021.

**Extended Data Fig 2:**
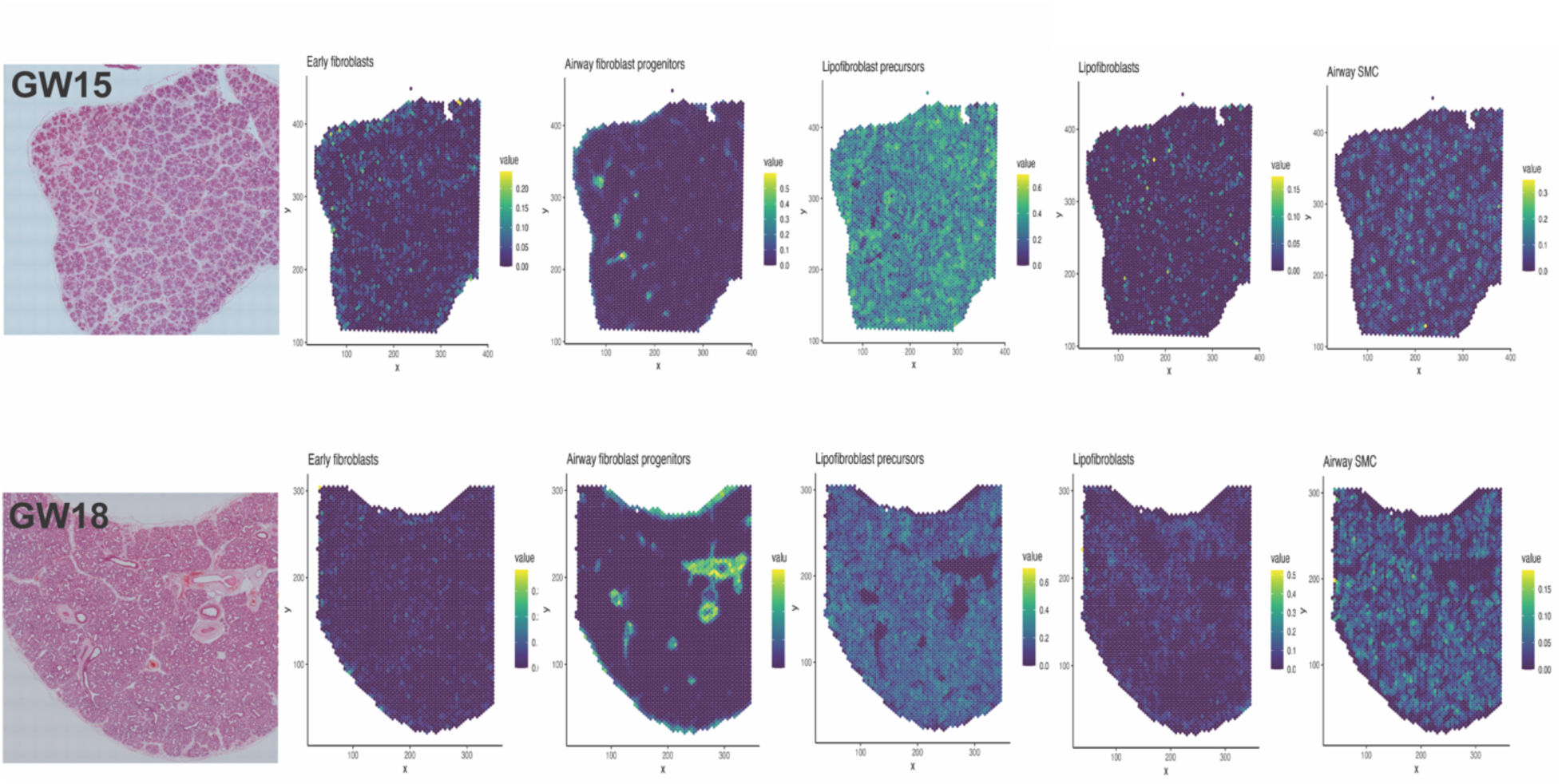
**Spatial transcriptomic of the fetal stromal cell compartment.** H&E staining shows the gross morphological features of the lung tissue. Visium spots with highest RCTD weights (green-yellow hue) for the major immune cell types. Top row: Representative GW15 fetal lung tissue. Bottom row: Representative GW18 fetal lung tissue.

**Extended Data Fig 3:**
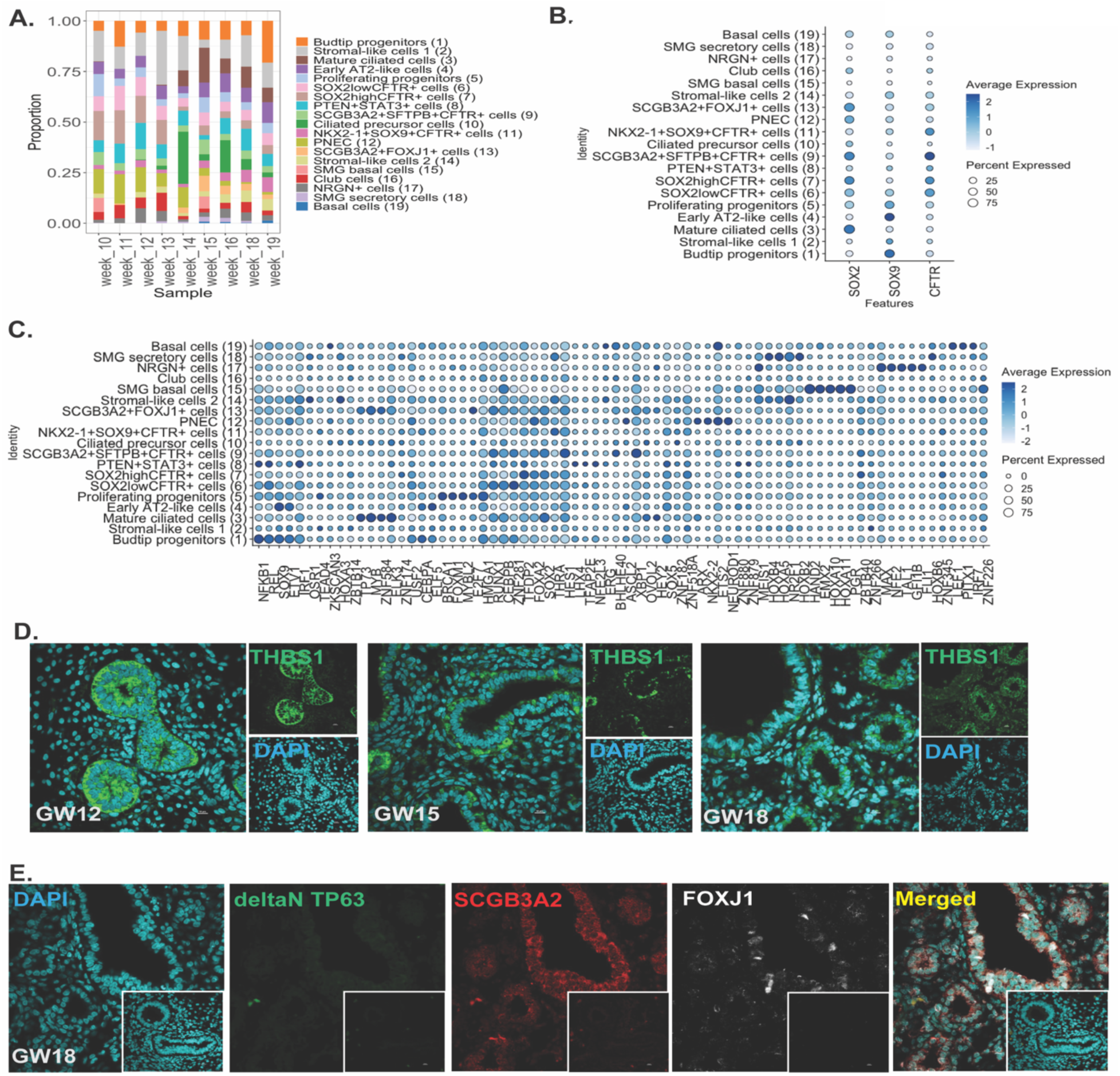
Characterization of the developing fetal lung epithelium. **A:** Proportion of the epithelial cell subtypes across GW. **B:** Dotplot for SOX2, SOX9 and CFTR to determine proximal, distal and cell expressing high levels of CFTR, respectively. **C:** Dotplot of the top differentially expressed transcription factors (TF) genes based on regulon specificity score (RSS) via *SCENIC*. **D:** Immunofluorescent staining THBS1, expressed in *PTEN+STAT3+* cells. Inset is the negative (no primary, secondary antibodies only) controls. DAPI marks all nuclei. **E:** Immunofluorescent staining deltaN P63 (basal cell), SCGB3A2 (secretory), and FOXJ1 (ciliated cell) in GW15 fetal lung epithelium. Inset is the negative (no primary, secondary antibodies only) controls. DAPI marks all nuclei.

**Extended Data Fig 4:**
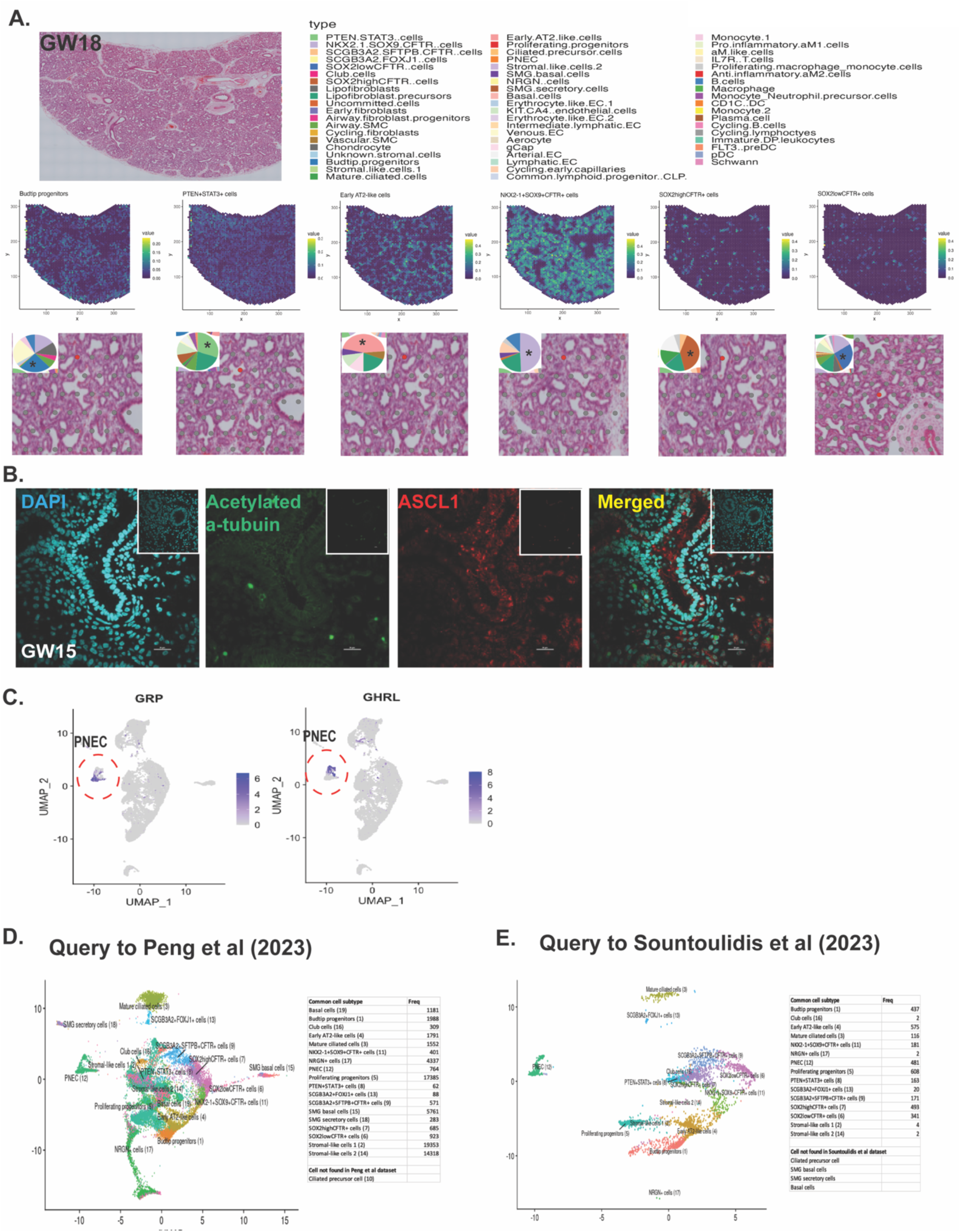
Spatial determination of epithelial subtypes in the fetal lungs and comparisons with published fetal lung datasets. **A:** Spatial transcriptomic analysis (10X Visium) shows areas of localization of budtip progenitors, *PTEN+STAT3+* cells, early AT2-like cells, *NKX2-1+SOX9+CFTR+* cells, *SOX2^low^CFTR+* cells, *SOX2^high^CFTR+* cells in GW18 fetal lung tissue. Location of Visium spots with highest RCTD weights (green-yellow hue) for the major immune cell types. High magnification of H&E staining shows a specific region where there is an abundant number of the specific cell subtype (red dot). Pie chart (insert) shows the proportion of the cells in the specific region (red dot). **B:** Immunofluorescent staining acetylated alpha tubulin (cilia on multiciliated cells) and ASCL1 (PNEC) in GW15 fetal lung epithelium. Inset is the negative (no primary, secondary antibodies only) controls. DAPI marks all nuclei. **C:** UMAP feature plots for *GRP* and *GHRL* demarcating two PNEC populations also identified in previous datasets. **D:** *MapQuery* analysis using our dataset as the reference map to identify shared cell subtypes with the dataset from Peng et al., 2022. **E:** *MapQuery* analysis using our dataset as the reference map to identify shared cell subtypes with the dataset from Sountoulidis et al., 2022.

**Extended Data Fig 5:**
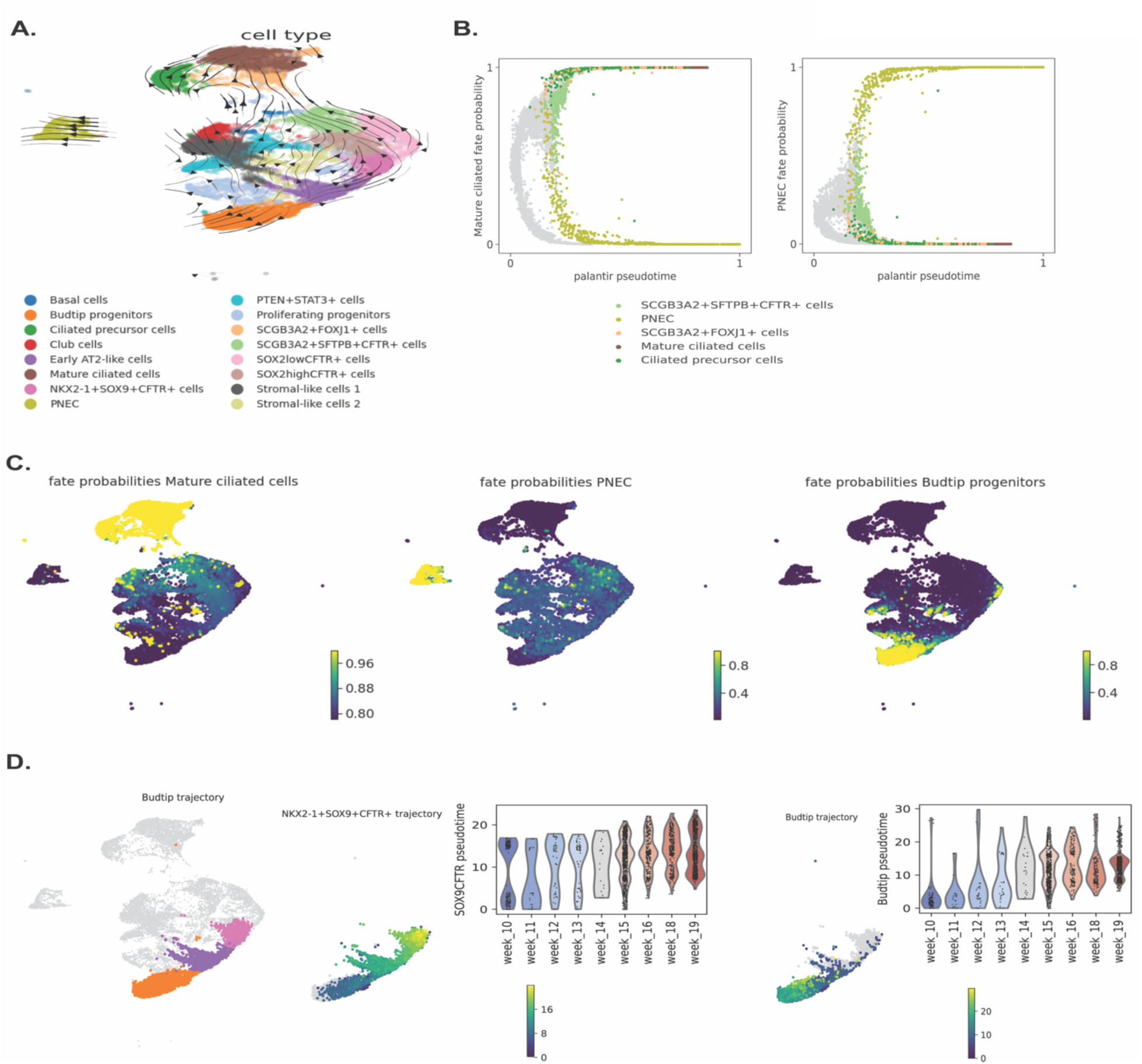
Validation of LatentVelo trajectories and analysis of distal trajectories. **A:** Palantir was used to infer pseudotimes, starting with a root at GW10 budtip cells. CellRank was used to infer a transition matrix between cells, and transitions were projected onto the UMAP. Results largely agree with LatentVelo, except for a clearer distinction of the terminal budtip state. The same terminal states at PNEC, Mature ciliated cells, and later GW budtip cells were observed. **B:** Mature ciliated cell and PNEC fate probabilities vs Palantir pseudotime. *SCGB3A2+SFTPB+CFTR+* cells were situated in the middle near 0.5, indicating their involvement in both lineages. **C:** Fate probabilities for the 3 terminal states. Fate probabilities for both the Stromal-like cells 1 and club cells, and *SCGB3A+SFTPB+CFTR+* and *SOX2+CFTR+* cells were elevated for the Mature ciliated and PNEC terminal states, as was also found with LatentVelo. **D:** Analysis of the budtip terminal state with Slingshot. A root was selected at early GW cells. Slingshot identified a bifurcation going towards the late GW budtip terminal state, or towards Early AT2-like cells and *NKX2-1+SOX9+CFTR+* cells. The Slingshot pseudotimes were correlated with GW.

**Extended Data Fig 6:**
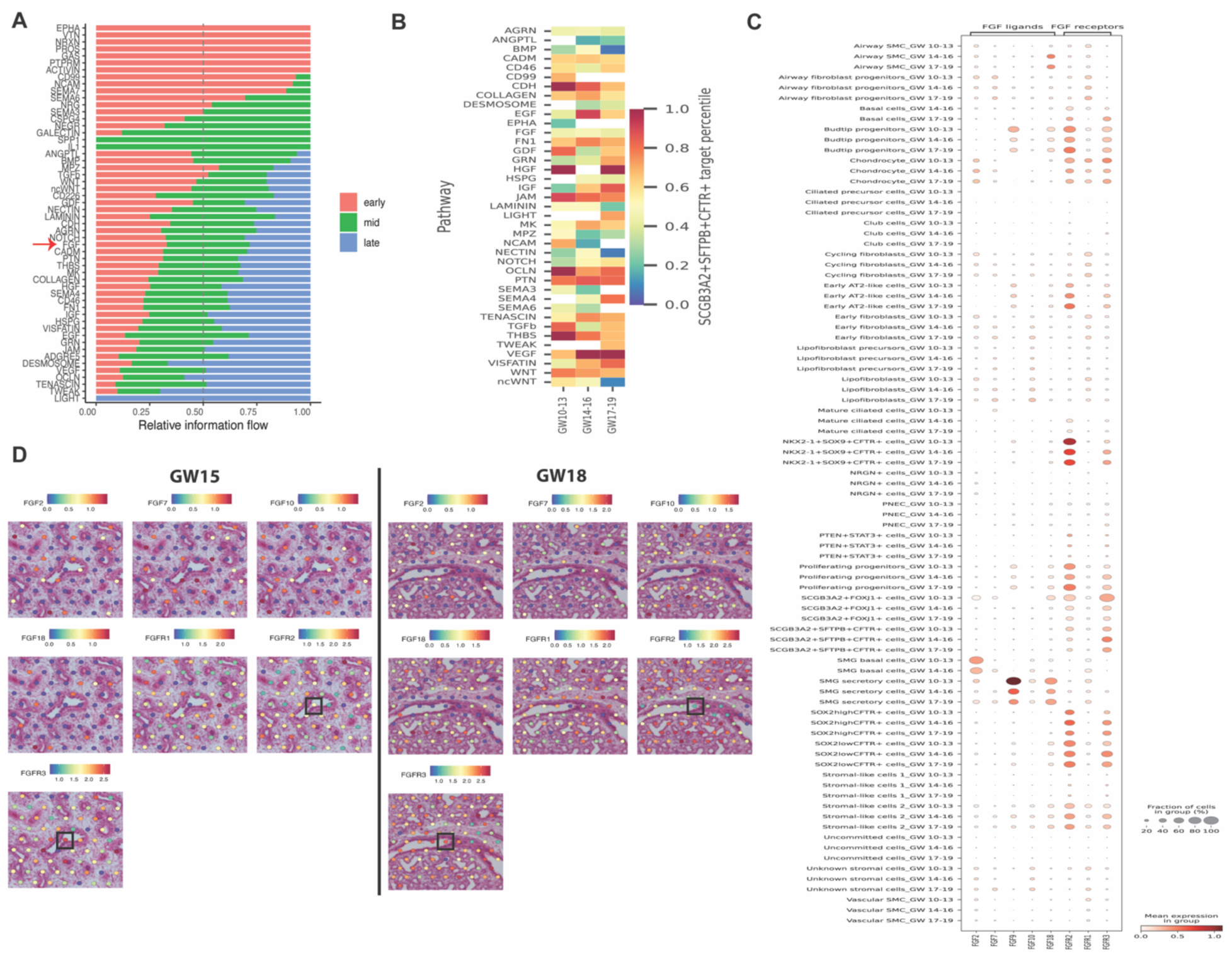
FGF cell signaling networks from senders to *SCGB3A2+SFTPB+CFTR+* cells. **A:** Relative information flow of signalling towards *SCGB3A2+SFTPB+CFTR+* cells for Early (GW10-13), Mid (GW14-16), and Late (GW17-19). Information flow is calculated as the sum of all cell interaction probabilities towards *SCGB3A2+SFTPB+CFTR+* cells. **B:** Overview of signaling pathways strength towards *SCGB3A2+SFTPB+CFTR+* cells in early (GW 10-13), mid (GW 14-16), late (GW 17-19) **C.** Expression of FGF ligands and receptors in specific epithelial and stromal subset of cells from scRNAseq grouped by early (GW 10-13), mid (GW 14-16), and late (GW17-19) stages. **D.** Expression of FGF ligands and receptors near *SCGB3A2+SFTPB+CFTR+* cells (black box on H&E staining) using Visium spatial transcriptomics.

**Extended Data Fig 7:**
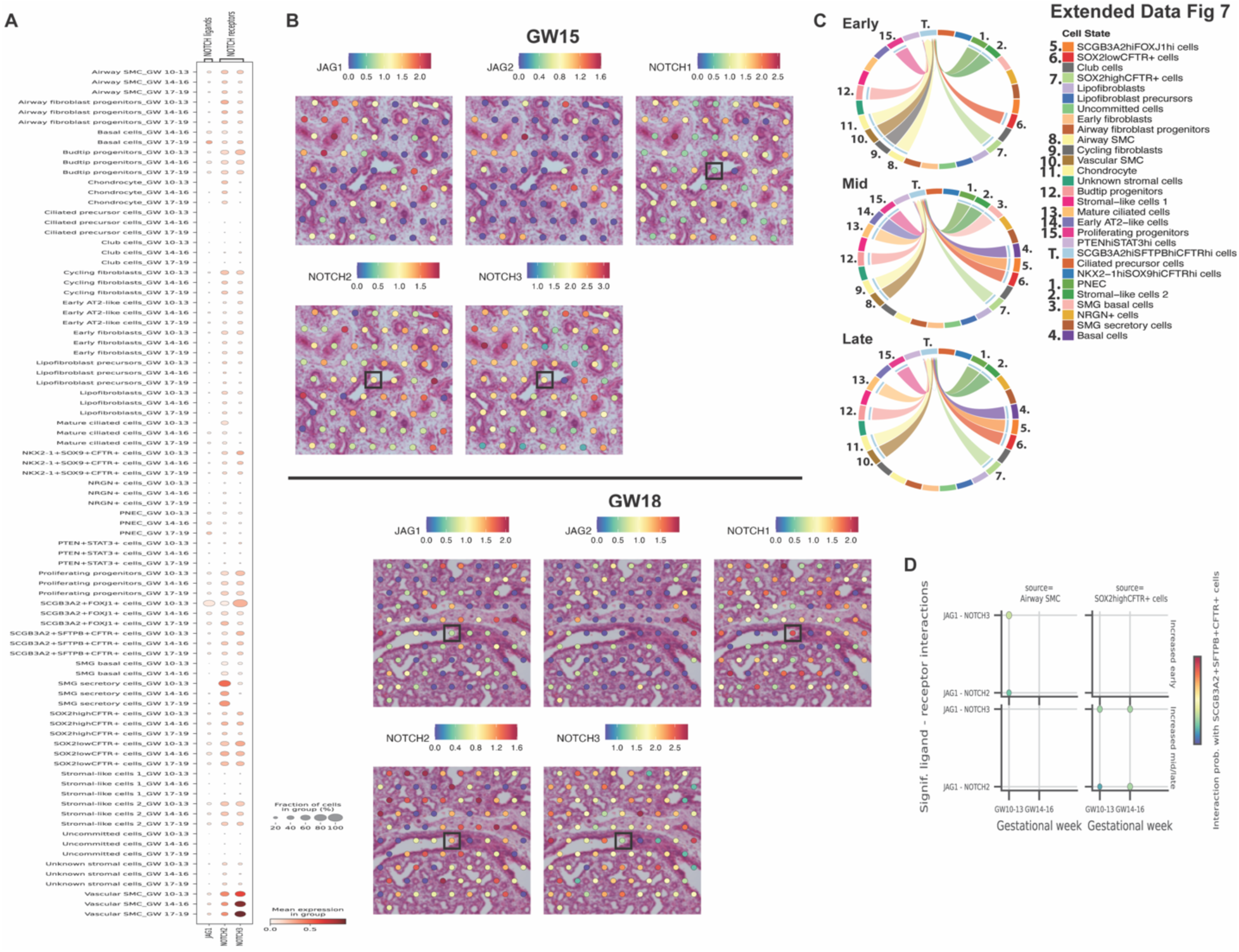
Notch signalling from senders to *SCGB3A2+SFTPB+CFTR+* cells. **A.** Expression of Notch ligands and receptors in specific epithelial and stromal subset of cells from scRNAseq grouped by early (GW 10-13), mid (GW 14-16), and late (GW17-19) stages. **B.** Expression of Notch ligands and receptors near *SCGB3A2+SFTPB+CFTR+* cells (black box on H&E staining) using Visium spatial transcriptomics. **C.** Chord diagrams demonstrating epithelial and stromal cell types that are senders of Notch signaling to *SCGB3A2+SFTPB+CFTR+* cells in early, mid, late stages. **D.** Ligand-receptor (L-R) plot showcasing specific NOTCH ligand-receptor interactions enriched in early vs mid/late from airway SMC, lipofibroblast precursors, airway fibroblast progenitors, and SMG secretory cells to *SCGB3A2+SFTPB+CFTR+* cells. Top show interactions significantly increased early, bottom shows interactions significantly increased mid/late. All tests use p < 0.01.

**Extended Data Fig 8:**
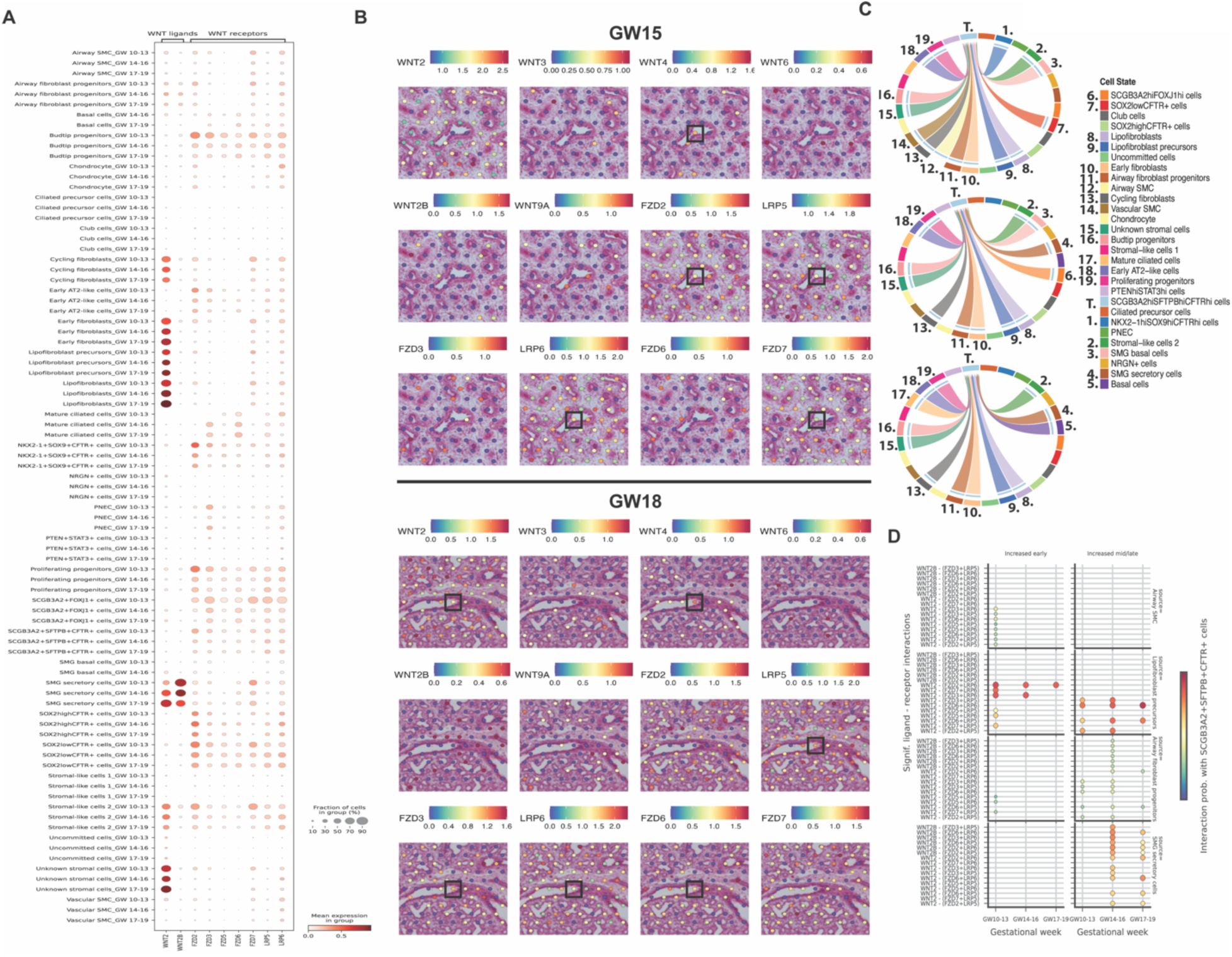
WNT signalling from senders to *SCGB3A2+SFTPB+CFTR+* cells. **A.** Expression of WNT ligands and receptors in specific epithelial and stromal subset of cells from scRNAseq grouped by early (GW 10-13), mid (GW 14-16), and late (GW17-19) stages. **B.** Expression of WNT ligands and receptors near *SCGB3A2+SFTPB+CFTR*+ cells (black box on H&E staining) using Visium spatial transcriptomics. **C.** Chord diagrams demonstrating epithelial and stromal cell types that are senders of WNT signaling to *SCGB3A2+SFTPB+CFTR+* cells in early, mid, late stages. **D.** Ligand-receptor (L-R) plot showcasing specific WNT ligand-receptor interactions enriched in early vs mid/late from airway SMC, lipofibroblast precursors, airway fibroblast progenitors, and SMG secretory cells to *SCGB3A2+SFTPB+CFTR+* cells. Top show interactions significantly increased early, bottom shows interactions significantly increased mid/late. All tests use p < 0.01.

## Supplementary Tables

**Table 1:** Detailed list of fetal lung tissue samples sequenced.

| **Sample ID**<br>**(Gestational week + day)** | **Sex** | **# of sequenced cells** | **# of genes** | **Filtered # of cells** | **% change of cells** |
| --- | --- | --- | --- | --- | --- |
| GW_10_1 | Female | 13852 | 27559 | 12412 | 0.103956107 |
| GW_10_2 | Male | 5937 | 17271 | 5515 | 0.07107967 |
| GW_11_1 | Male | 4923 | 16935 | 4629 | 0.059719683 |
| GW_12_3 | Male | 10689 | 17067 | 10277 | 0.038544298 |
| GW_13_4 | Male | 3748 | 16573 | 3588 | 0.042689434 |
| GW_13_6 | Male | 13464 | 17276 | 13054 | 0.030451575 |
| GW_14_3 | Male | 3916 | 16255 | 3748 | 0.042900919 |
| GW_15_5 | Female | 15444 | 18067 | 8168 | 0.471121471 |
| GW_15_5_1 | Female | 8849 | 17214 | 13347 | -0.508306023 |
| GW_16_1 | Male | 6425 | 16835 | 5636 | 0.122801556 |
| GW_16_2 | Female | 7687 | 16415 | 7380 | 0.039937557 |
| GW_16_5 | Female | 5524 | 24481 | 4566 | 0.173425054 |
| GW_18 | Male | 10505 | 16700 | 10477 | 0.002665397 |
| GW_18_1 | Male | 12109 | 16855 | 10876 | 0.101825089 |
| GW_18_2 | Female | 6330 | 16223 | 6060 | 0.042654028 |
| *GW_18_1A | Male | 7621 | 17182 | 6920 | 0.091982679 |
| *GW_18_1B | Male | 6341 | 17061 | 5888 | 0.071439836 |
| GW_19_0 | Male | 10565 | 17664 | 9764 | 0.075816375 |
| GW_19_2 | Female | 6847 | 16970 | 6107 | 0.10807653 |
| GW_19_4 | Male | 9480 | 25027 | 8286 | 0.125949367 |
| **Total count** | **7 Female**<br>**12 Male** | **170256** |  | **156698** |  |

\*Same gestational lung tissue. “A” and “B” indicates different regions (lower and upper regions) of the lung tissue sampled for sequencing.

**Table 2:** List of cell type markers top differentially expressed genes. Please see attached excel file.

**Table 3:**
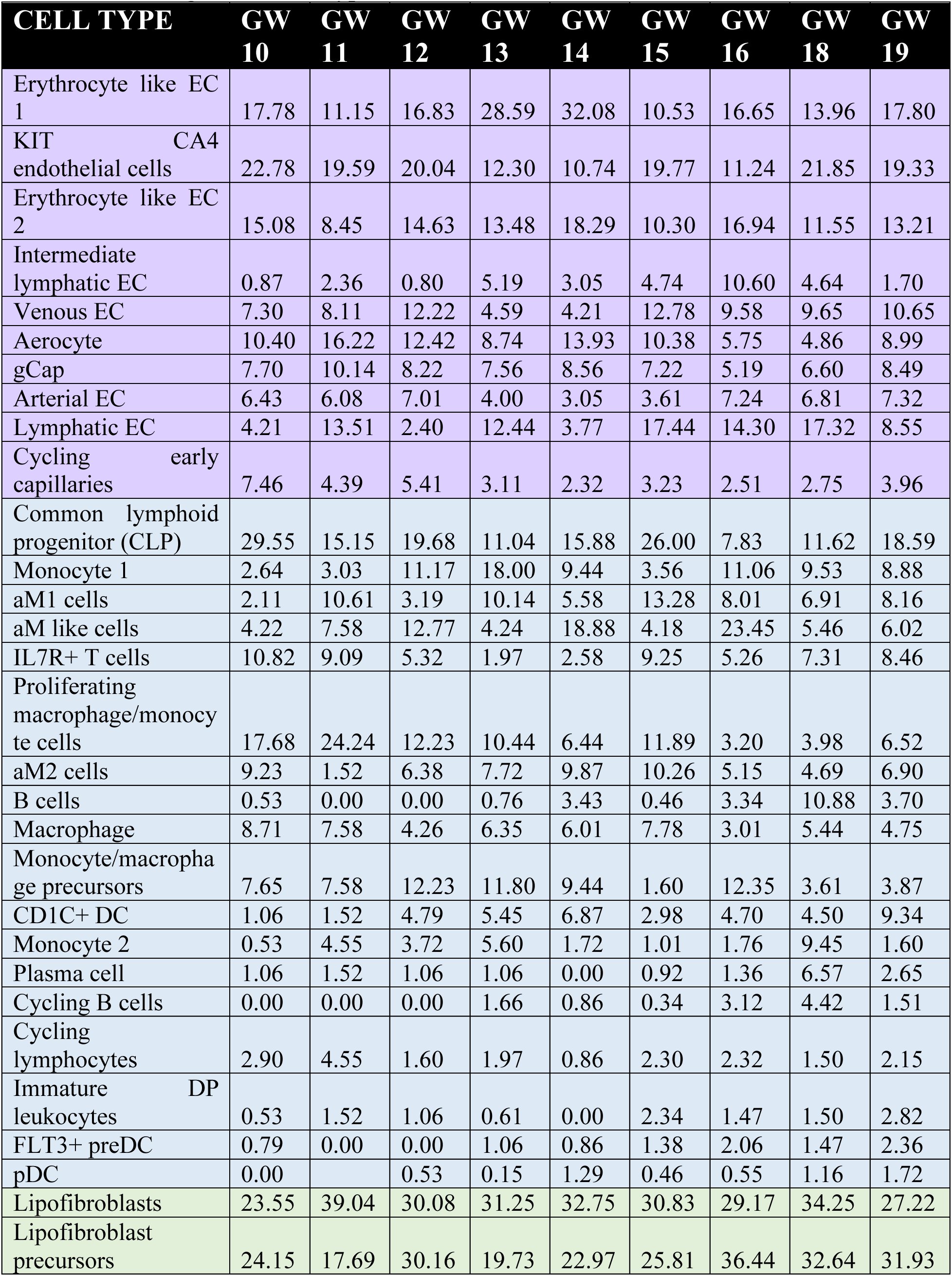
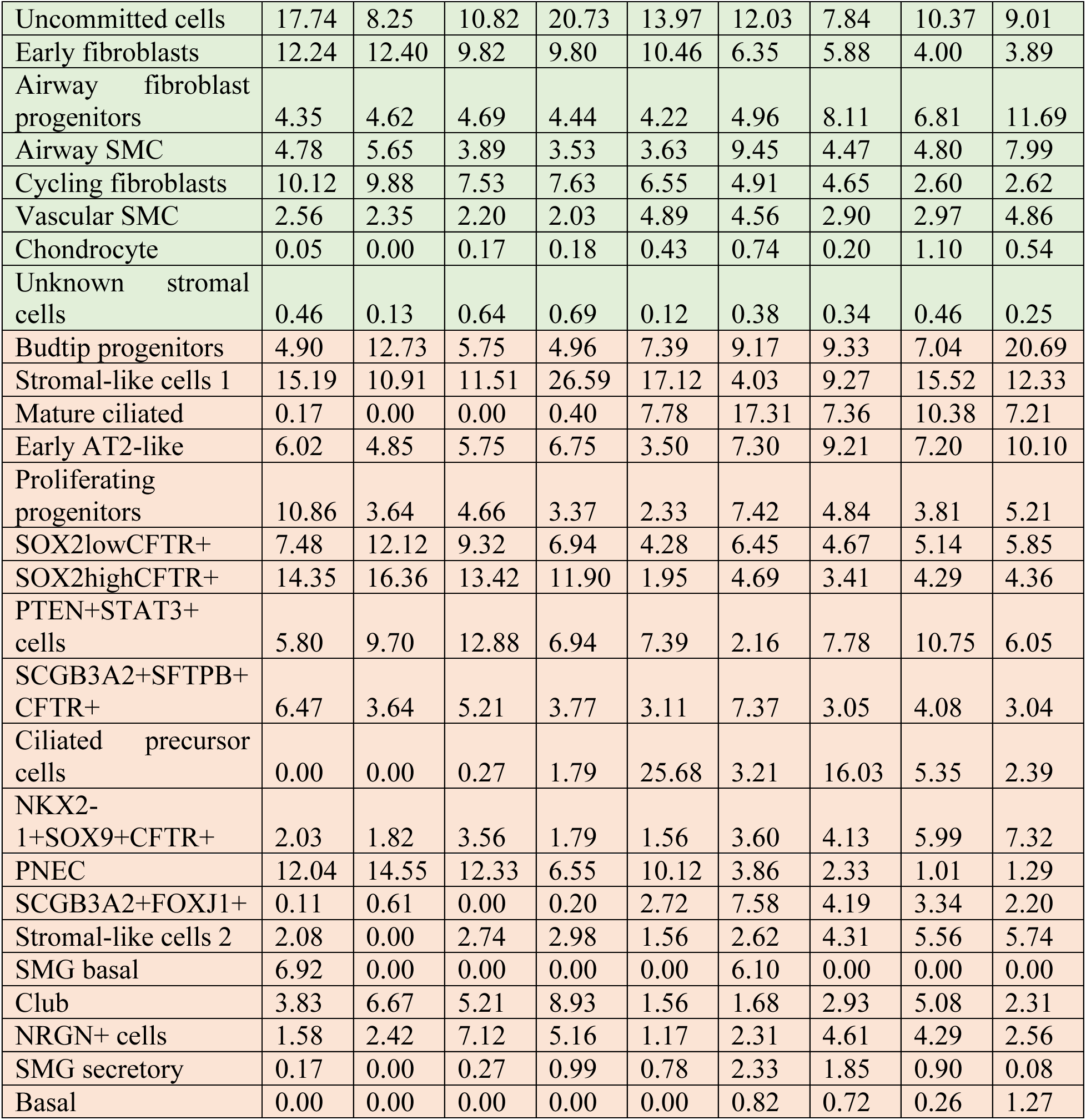
Percentages of cell subtypes.

**Table 4:** List of primary and secondary antibodies.

| **Primary Antibodies** | **Host** | **Supplier** | **Cat. no.** | **Dilution** | **Assay** |
| --- | --- | --- | --- | --- | --- |
| Acetyl-alpha tubulin | Mouse | Invitrogen | 322700 | 1:200 | IF |
| ASCL1 | Rabbit | Abcam | Ab74056 | 1:200 | IF |
| CFTR | Mouse | Millipore | MAB1660 | 1:70 | IF |
| CGRP | Mouse | Sigma | C9487 | 1:100 | IF |
| Delta NP63 | Rabbit | Biolegend | 619002 | 1:200 | IF |
| FOXJ1 | Rabbit | Sigma | HPA005714 | 1:175 | IF |
| SOX2 | Goat | R&D | AF2018 | 1:300 | IF |
| SOX9 | Goat | R&D | AF3075 | 1:600 | IF |
| SCGB1A1 | Rat | R&D | MAB4218 | 1:50 | IF |
| SCGB3A2 | Goat | Santa Cruz | sc-48320 | 1:50 | IF |
| PGP9.5 | Mouse | Abcam | ab20559 | 1:100 | IF |
| ProSPB | Rabbit | Abcam | ab40876 | 1:50 | IF |
| ProSPC | Rabbit | Abcam | ab40879 | 1:100 | IF |
| THBS1 | Mouse | LS Bio | LS-C87 | 1:50 | IF |
| TTF1 | Rabbit | Abcam | ab76013 | 1:200 | IF |

| **Secondary Antibodies** | **Host** | **Supplier** | **Cat. no.** | **Dilution** | **Assay** |
| --- | --- | --- | --- | --- | --- |
| Anti-rabbit IgG (H+L) Alexa Fluor 488 | Donkey | Invitrogen | A21206 | 1:500 | IF |
| Anti-mouse IgG (H+L) Alexa Fluor 546 | Donkey | Invitrogen | A10036 | 1:500 | IF |
| Anti-goat IgG (H+L) Alexa Fluor 647 | Donkey | Invitrogen | A21447 | 1:500 | IF |
| Anti-mouse IgG (H+L) Alexa Fluor 488 | Donkey | Invitrogen | A21202 | 1:500 | IF |
| Anti-goat IgG (H+L) Alexa Fluor 546 | Donkey | Invitrogen | A11056 | 1:500 | IF |
| Anti-Rabbit IgG (H&L) Alexa Fluor 647 | Donkey | Abcam | ab150075 | 1:500 | IF |
| Anti-rat IgG (H+L) Alexa Fluor 555 | Donkey | Invitrogen | A48270 | 1:500 | IF |
| Anti-rabbit IgG (H+L) Alexa Fluor 546 | Donkey | Invitrogen | A10040 | 1:500 | IF |

## Supplementary Note 1

### Co-development of the fetal pulmonary endothelium

The fetal pulmonary endothelium contains 10 endothelial subtypes in the early developing lung (**Suppl. Note Fig. 1A**) that expressed pan-endothelial genes *PECAM1*. These endothelial cell (EC) subtypes include erythrocyte-like EC-1, *KIT+CA4+* EC, erythrocyte-like EC-2, intermediate lymphatic EC, lymphatic EC, venous EC, aerocytes, general capillaries (gCap), arterial EC, and cycling early capillaries. Both EC-1/EC-2 expressed high levels of hemoglobin genes and were annotated as erythrocyte-like cell populations (**Supplementary Table 2**). The proportion of each EC subtypes remained relatively consistent in all the gestational timepoint studied (**Suppl. Note Fig. 1B**). Notably, the fetal lung lymphatic cells were found as early as GW10 and continued to develop with time, which would become important in neonatal lung respiration^1^. The top DEG expressed in both lymphatic populations included *PROX1*, *CCL21* and *TFF3*^2,3^ with the intermediate lymphatic EC expressed higher levels of genes associated with inflammation *PTX3* and *NRP2* (**Suppl. Note Fig. 1C** and **Supplementary Table 2**). Differential enrichment analysis of the transcription factors driving these cell lineages was performed using SCENIC^4^ (**Suppl. Note Fig. 1D**). Cycling early capillaries expressed high levels of genes associated with cellular proliferation such as *TOP2A, MKI67*, and *CENPF.* Not surprisingly, transcription factors regulating cell proliferation and cell cycle *BRCA1, E2F7, MYBL2* were highly expressed in the cycling early capillaries. Aerocytes and gCap cells shared similar DEG with gCap expressing higher levels of *HBEGF,* previously shown to promote angiogenesis^5^, while aerocytes differentially expressed high levels of *EGFL6* and *COL6A3*. Interestingly aerocytes share similar differentially expressed transcription factors as gCap, including *HOXB4, IRF, POU2F2,* suggesting these two cell types may share developmental origins, as has been previously shown in adult mouse lungs^6^. Higher expression of *HGPD, CA4, KIT* distinguished the *KIT+CA4+* cells from the EC-1 and EC-2. Higher expression of the transcription factors *MLX, SOX18* and *ZNF71* were also observed in KIT+CA4+ EC. Venous EC expressed high levels of the specific pulmonary venous gene *CPE*^3^ and the apelin receptor *APLNR*, the latter previously shown to regulate endothelial cell differentiation^7^. Arterial EC expressed abundant *DKK2*, a Wnt pathway modulator and the chemokine ligand *CXCL12* and the transcription factors *CREB1* and *SMAD4,* both important in the formation and function of the vasculature^8,9^.

To determine the developmental relationships between these cells, we used our RNA velocity-based method *LatentVelo*^10^ (**Suppl. Note Fig. 1E and E’**, respectively) and visualized velocities using partition-based graph abstraction (PAGA) of RNA velocity projections (**Suppl. Note Fig. 1F**). In brief, velocities infer the directionality of lineage development and corresponding latent times identify the position of cells along the developmental trajectory, shown as a dark blue to yellow hue (**Suppl. Note Fig. 1E’)**. In doing so, cell origins were determined, and trajectories predicted based on these relationships over time. Several conserved trajectories were found which included unique trajectories stemming from putative multipotent EC. Focusing on capillary specification, we identified a lineage relationship between cycling early capillaries to gCap cells and aerocytes with a gradual change in top DEGs associated with these cell lineages as they differentiate (increase in *CXCL12,* and *EDNRB* associated with gCap and aerocyte differentiation respectively, **Suppl. Note Fig. 1G**). Interestingly, this same lineage development of gCap and aerocytes has also recently been found in adult distal lungs^6^. Spatial transcriptomics of GW15 fetal lung showed uniform distribution of both gCap and aerocytes throughout the tissues (**Suppl. Note Fig. 1H**).

### Novel alveolar macrophage lineage trajectories identified in the fetal lung

The fetal lung immune population contains 18 distinct cell types (**Suppl. Note Fig. 2A**). These include a common lymphoid progenitor, proliferating macrophage/monocytes, monocyte 1 and 2, anti-inflammatory alveolar macrophage 2 (aM2), proinflammatory aM1, aM-like cell, B cells, cycling B cells, plasma cells, monocyte/neutrophil precursor cells, *CD1C+* dendritic cells 1 (DC), macrophage, *FLT3*+ preDC, *IL7R+* T cells, immature double positive (DP) leukocytes, cycling lymphocytes, and plasmacytoid DC (pDC). The proportion of mature cell types including monocytes, plasma cells and B cells increased in later gestational lung tissues (**Suppl. Note Fig. 2B**). Top genes differentially expressed in each subtype distinctly separates the different clusters (**Suppl. Note Fig. 2C** and **Supplementary Table 2**). Gene ontology enrichment analysis of the DEG (**Suppl. Note Fig. 2D**) showed terms associated with antigen processing and presentation and major histocompatibility complex (MHC) class II protein in all DC lineages. Dendritic cells made up a smaller proportion of myeloid cells in the fetal lungs. Their role in immunity during development is unknown. However, studies have suggested fetal DC may help induce immunologic tolerance where fetal and maternal immune cells may come into contact during pregnancy^12^. *CD1C+* DC and *FLT3+* DC share several common genes mainly the HLA family genes. *CD1C+* DC have previously been shown to prime cytotoxic T cell responses^13^, while *FLT3+* DC may mark an early developing DC population as the FLT3 ligand has previously been shown to regulate dendritic cell development^14^. Transcription factors differentially enriched (**Suppl. Note Fig. 2E**) in each cell cluster identified unique expression of acetyl-CoA carboxylase 1 (*ACAA1)* in *CD1C+* DC and Basic Leucine Zipper ATF-Like Transcription Factor 3 *(BATF3)* in *FLT3+* DC, both previously shown to regulate DC development and function^15,16^, respectively. Spatial transcriptomic show both *CD1C+* and *FLT3+* DCs sparsely distributed in the fetal lung tissue (**Suppl. Note Fig. 2F**).

The AT-rich interaction domain *3A (ARID3A)* transcription factor was differentially enriched in pDC, as previously observed^17^. Plasmacytoid DC are rare cells that secrete large amounts of type I interferons with a role in antiviral immunity and share similar morphology as plasma cells^18^. They also play a role in inducing T cell tolerance and therefore may exist in the developing lung to suppress alloreactive T cells. While this is the first identification of pDC in the fetal lung, it is unclear if these pDC originated in the developing lungs or as previous studies have shown, are bone marrow-derived and circulated into the lungs^19^, but play an important role in immune responses to inhaled antigens^20^. LatentVelo identified a lineage trajectory originating from pDC to *CD1C+* DC to aM2 cells and aM1 cells, and a trajectory from *FLT3+* preDC to *CD1C+* DC (**Suppl. Note Fig. 2G-H**). GO terms associated with inflammatory and immune responses were enriched in aM1, aM2 and AM-like clusters (**Suppl. Note Fig. 2D**). Both aM1/aM2 are delineated based on *CD68* expression, a classical marker of monocyte-derived alveolar macrophages, and were distinguished between one another based on expression of proinflammatory genes *IL1B, TNF* and anti-inflammatory *IL10* and *TGFB* genes. On the contrary, aM-like cells expressed *IL1B* but shared no other proinflammatory genes as aM1. Alveolar macrophages (aM-like, aM1, aM2) shared similar elevated expression of *POU2F2* and *NFKB1* suggesting a common conserved developmental regulation (**Suppl. Note Fig. 2E**). Regulation of *NFKB1* signalling in fetal macrophages is important for airway branching as it protects from LPS-induced proinflammatory responses^21,22^. High expression of *SMAD6* distinguished aM-like from aM1 and aM2 cells, while *KLF11* expression was higher in aM2 cells. Moreover, aM-like cells expressed *IL1B* but shared no other proinflammatory genes with aM1. Spatially, aM-like, aM1 and aM2 cells were found sparsely distributed aM1 and aM2 cells (**Suppl. Note Fig. 2F**). LatentVelo and Slingshot analyses (**Suppl. Note Fig. 2G, G’**) showed a lineage trajectory originating from monocyte/macrophage precursor to aM-like cells to aM2 cells and aM1 cells which is confirmed with a corresponding change in top DEG as the cells differentiate along the trajectories (**Suppl. Note Fig. 2H and I**).

B cells, cycling B cells, and plasma cells express the canonical B cell signalling molecules *CD79*, *VPREB3*^23^ and GO terms associated with B cell receptor signaling pathway and B cell activation (**Suppl. Note Fig. 2D**). Indeed, bacterial LPS is a potent stimulant of B cell and macrophage differentiation and antibody secretion^24^. LatentVelo and Slingshot analyses identified a lineage trajectory originating from B cells to plasma cells, and trajectories from monocyte/macrophage precursor to monocyte 1 and monocyte 2 (**Suppl. Note Fig. 2I**).

**Supplementary Note Figure. 1:**
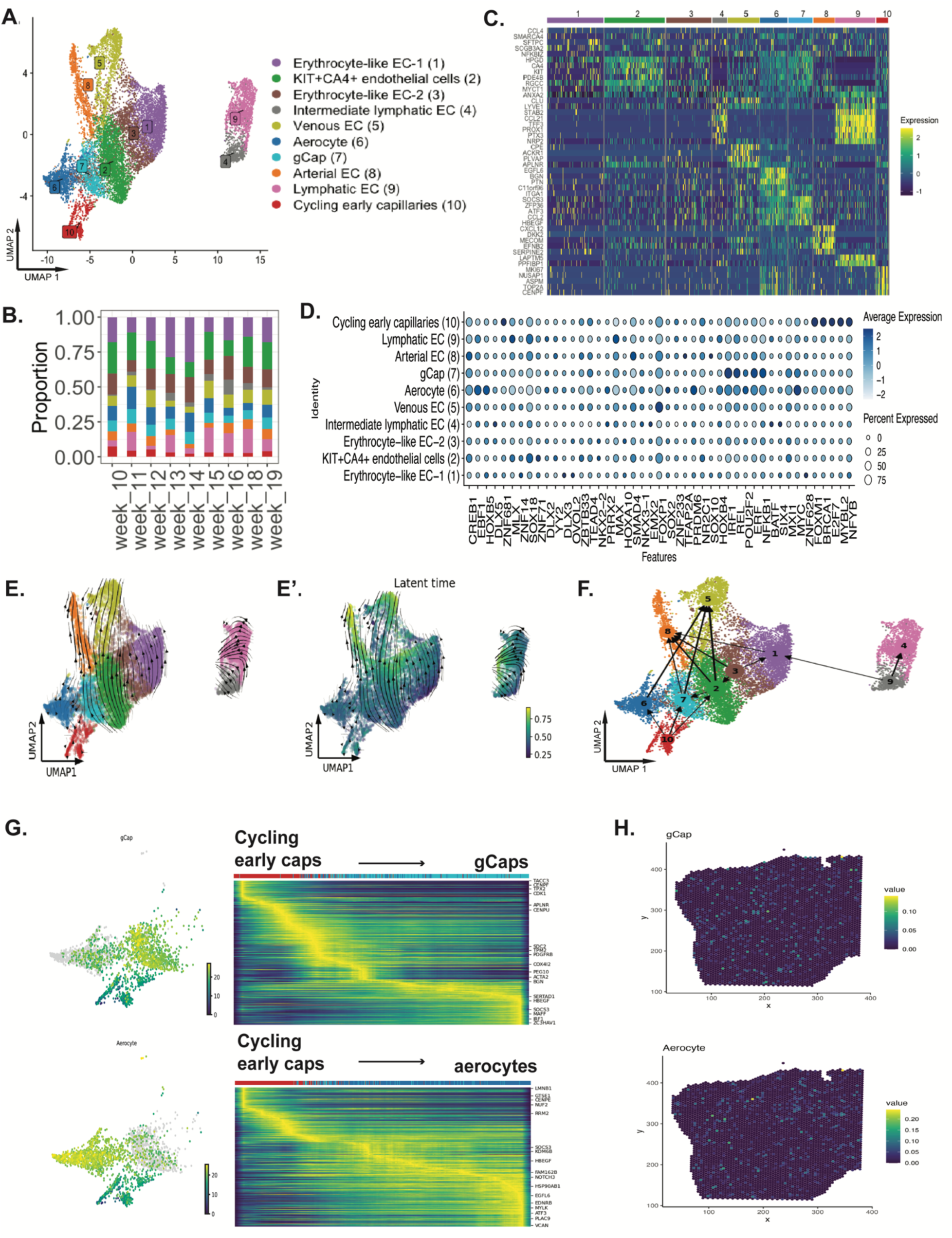
Co-development of the fetal pulmonary endothelium. **A:** UMAP visualization of the fetal endothelial subtypes. **B:** Gene expression heatmap of the top 5 differentially expressed genes in each cluster. **C:** Proportion of the endothelial cell types across GW. **D:** Dotplot of the top differentially expressed transcription factors (TF) genes based on regulon specificity score (RSS) via *SCENIC*. **E:** UMAP projection of inferred LatentVelo velocities and **E’:** LatentVelo latent times. **F:** PAGA using LatentVelo velocities. **G:** Slingshot trajectory analysis using a root at Cycling early capillaries, as identified by LatentVelo. Trajectories to gCaps and Aerocytes are identified. Trajectory heatmaps indicate the progression of celltypes and significantly varying genes along the trajectories. **H:** Location of Visium spots with highest RCTD weights (green-yellow hue) for gCaps and Aerocytes.

**Supplementary Note Figure. 2:**
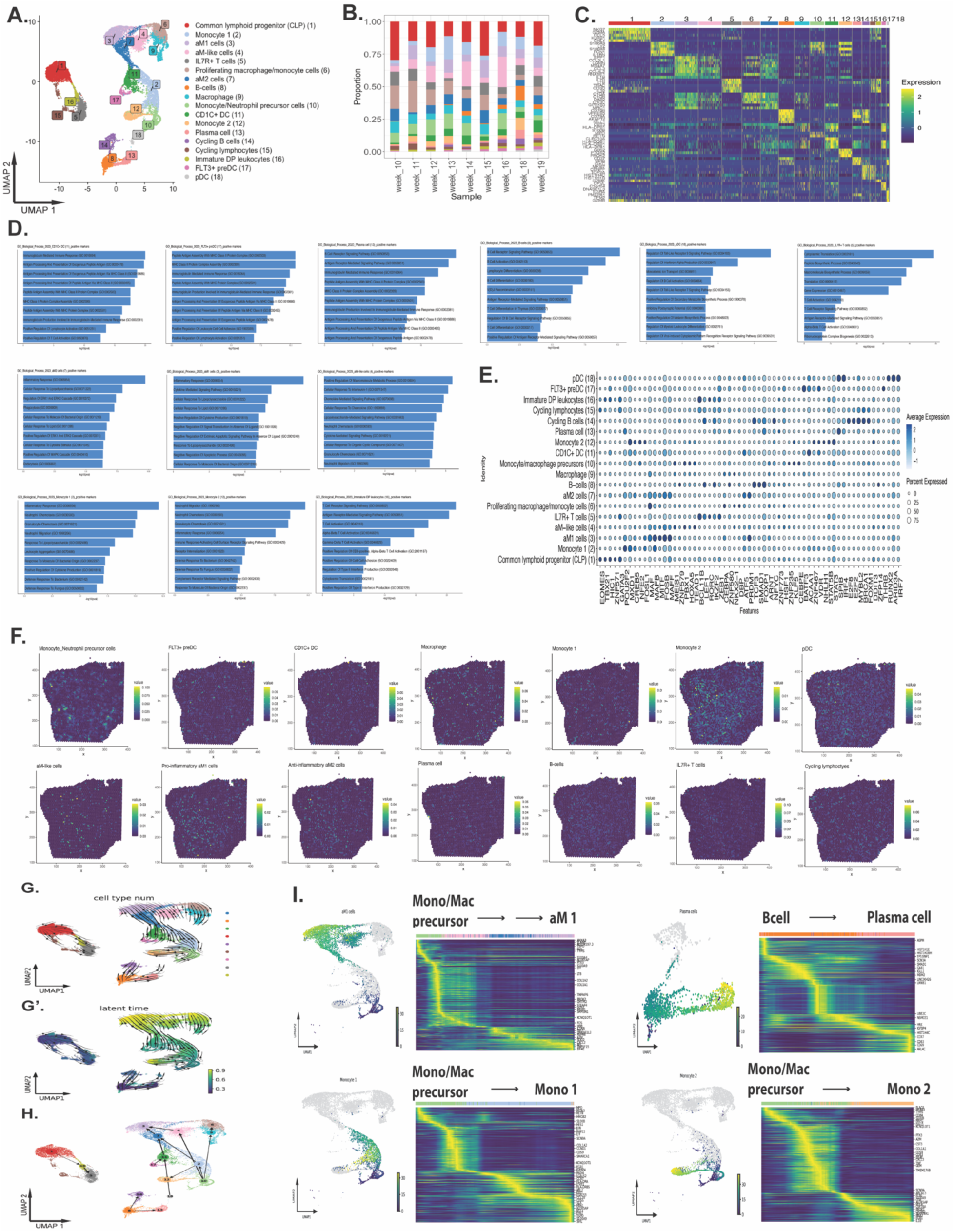
Novel alveolar macrophage lineage trajectories identified in the fetal lung. **A:** UMAP visualization of the fetal immune cell subtypes. **B:** Proportion of the immune cell types across GW. **C:** Gene expression heatmap of the top 5 differentially expressed genes in each cluster. **D:** Gene ontology terms enriched each of the major immune cell subtypes. **E:** Dotplot of the top differentially expressed transcription factors (TF) genes based on regulon specificity score (RSS) via *SCENIC*. **F:** Location of Visium spots with highest RCTD weights (green-yellow hue) for the major immune cell types. **G:** UMAP projection of inferred LatentVelo velocities and **G’:** LatentVelo latent times. **H:** PAGA using LatentVelo velocities. **I:** Slingshot trajectory analysis using a root at Monocyte/Macrophage precursor cells, as identified by LatentVelo. Trajectories to aM1, and Monocyte 1 and 2 are identified. Using a root at B cells, a trajectory is inferred to Plasma cells. Trajectory heatmaps indicate the progression of celltypes and significantly varying genes along the trajectories.

## Materials & Methods

### Human fetal lung collection

Human fetal lung tissues were collected by the Research Centre for Women’s and Infants’ Health (RCWIH) biobank. All tissue collections were approved by the Mount Sinai Hospital Research Ethics Board and the Hospital for Sick Children Research Ethics Board for collection and research use of gestational week lung tissues from 10-20 weeks. Freshly isolated human fetal lung tissues were collected via careful microdissection by an experienced nurse and stored in HBSS (Hank’ Balanced Salt Solution) on ice during the transfer from the biobank to the Hospital for Sick Children for immediate processing. Tissues were absent of respiratory abnormalities or known genetic lung defects.

### Sample preparation for library construction

The fetal lung samples were dissociated into single-cell suspension using the Multi Tissue Dissociation Kit 1 (Miltenyi Biotec; cat # 130-110-201) and 37C Multi_tissue_dissociation B program. Dissociated cells were collected through a 40 µm pore size cell strainer (Falcon; cat # 352340) and incubated with RBC lysis buffer (Invitrogen; cat# 00-4333-57) for two minutes to remove red blood cells. Cell number and viability were assessed by trypan blue staining (Gibco; cat# 15250061) and counted with the Countess II (Life Technologies, cat#A27977). This procedure resulted in cell viability of at least 80%. Approximately ∼10,000 cells per sample were captured and used for library construction using the 10X Chromium Next GEM single-cell 3’ Reagent Kits v3.1 (10x Genomics, cat#1000121, 1000120, 1000123). Library prep was performed as per manufacturer’s protocol. Sequencing was performed on the NovaSeq6000 (The Centre for Applied Genomics (TCAG) Sequencing Facility, SickKids). The target reads were 60,000 per cell and approximately 3,000-10,000 cells per sample were sequenced.

### Quality control and data processing

Raw FASTQ files were generated using *supernova/cellranger* mkfastq and bcl2fastq v2.20. 10x Genomics *Cell Ranger v6.0.1* software was used to align reads to the human reference genome (hg 19, GRCh38). *Seurat v4.0*^1^ was used for subsequent analysis. Cells with less than 200 features, greater than 15% mitochondrial transcript, and genes expressed in less than 3 cells were excluded from our analysis. Furthermore, principal component analysis (PCA) for each sample was done to ensure filtering of doublets and high-quality cells. Gene expression levels were log normalized datasets and highly variable features that identify high cell-cell variation within each sample dataset were determined using Seurat‘s *FindVariableFeatures*. Subsequently features that were repeatedly variable across each sample dataset was identified as integration features. Data was then scaled (linear transformation) was done using Seurat’s *ScaleData* and principal component analysis (PCA) (linear dimensional reduction) was done on each sample dataset. Generation of a master dataset was done by integrating all the sample datasets using Seurat’s reciprocal PCA (RPCA) integration (3000 features). Subsequent PCA analysis was done to ensure proper integration and 30 dimensions were retained for further analysis. Further reduction was done through uniform manifold approximation and projection (UMAP) (non-linear dimensional reduction) and cluster analysis was done using Seurat’s *FindClusters*. Biological sex was determined based on *SRY, XIST*, and *DDX3Y* expression.

### Clustering and differentially expressed genes analysis

The clustree package^2^ was used to inform the correct resolution for sub-clustering. Clustree assess the stability of the clusters generated by taking the overlap in cell clusters across multiple resolutions and calculating the in-proportion for each edge. Resolutions with clusters derived from high in-proportion score of multiple parent clusters were deemed over-clustered. Manual annotations were performed for each clusters and DEG compared to He et al. 2022 (courtesy of Dr. Emma Rawlins)^3^. Differentially expressed genes (DEGs) for each cluster were calculated using Seurat’s *FindAllMarkers*. Parameters that were defined included logFC > 0.25 and only output positive values. This analysis was done on the whole dataset where canonical markers identified within the top DEGs were used to assign main fetal cell type identities and within each cluster to determine DEGs in sub-clusters for sub-cluster analysis.

Cell-type scores for each cluster based on these DEGs are computing using the scanpy “sc.tl.score_genes” function. The top 100 DEGs for each cluster are used as the gene set, and all other genes expressed in more than 10 cells are used as the background reference pool.

### Gene ontology and transcription factor enrichment analysis

Gene ontology (GO) analysis was done using ‘DEenrichRPlot’ function accessing *EnrichR* databases: GO_Biological_Process_2023, GO_Cellular_Component_2023, and GO_Molecular_Function_2023 on top 100 DEG for each sub cluster^4^. Enrichment of top 10 GO terms (ordered by log p-value). DEGs were subsequently used for transcription factor enrichment analysis using SCENIC (pySCENIC version 0.11.2)^5^. First, gene regulatory interactions were calculated based on co-expression across the single-cell dataset with GRNBoost2^6^, followed by pruning interactions using known TF binding motifs and the construction of dataset specific regulatory modules (regulons)^7^. Regulons were then scored in each individual cell using AUCell and regulon specificity score (RSS) was subsequently calculated to identify enriched transcription factors in each cluster.

### Spatial transcriptomics

Spatial transcriptomics was performed using the 10X Visium platform (FFPE v2) and processed as per manufacturer’s protocol. For this, archived human fetal lung tissues (GW 15 and GW 18) from paraffin-embedded blocks were freshly sectioned, tested for RNA quality control (DV200 > 30%) before mounting on the 10X CytAssist (6.5 mm x 6.5mm) for library prep and sequencing (∼50,000 reads/spot). H&E staining was done as per manufacturer’s protocol. The Visium Human Transcriptome Probe Set v2.0 was used. Spaceranger (2.1.0) was used to perform demultiplexing and alignment to the GRCh38-2020-A reference. Spots with > 10% mitochondrial content was excluded from analysis. SCTransform normalization, PCA (30 principal components was used for downstream analysis), and UMAP was done in *Seurat* (v4.3.0.1)^1^. Robust cell-type decomposition was done using *spacxr* as published^8^. In brief, counts and annotations from scRNA-seq dataset was used as a reference and counts from Visium were used as a query. Spots were deconvolved using ‘full’ mode and RCTD weights for each spot were collected. High proportion of cells was determined using a threshold of 0.1.

### Trajectory analyses

Standard RNA velocity pre-processing is done using scVelo by normalizing cells by library size with scv.pp.filter_and_normalize, and filtering genes with less than 100 cells expressing unspliced and spliced counts for the gene with scv.pp.filter_genes (except in the epithelial where we use 30 and stromal where we use 300). The top 3000 highly variable genes are selected with scv.pp.filter_genes_dispersion with flavor=’cellranger’. Following the standard scVelo preprocessing, moments are computed by averaging over 100 nearest neighbors computed on 30 principle components from the log(1+x) transformed spliced counts with scv.pp.moments (300 nearest neighbors for stromal). We normalize each gene by its standard deviation, and input to LatentVelo for velocity inference.

We run LatentVelo for each population separately (stromal, epithelial, endothelial, and immune populations), utilizing the batch information from each of the 19 samples. We set the latent dimension of LatentVelo as 50 and encoder hidden layer size of 75 for the endothelial and immune populations, and 100 and 125 for the epithelial and stromal. The dimension of the latent regulatory state is changed according to the complexity of the dataset, and has dimension 2 for stromal, 4 for epithelial, 3 for endothelial, and 3 for immune. When analyzing the epithelial cells with LatentVelo, we remove of SMG secretory, SMG basal, and NRGN+ cells, since these small clusters were not strongly connected to any of the larger clusters. For the stromal cells, we subset to just the fibroblast populations.

We use standard scVelo functions to visualize velocities. To project latent velocities onto UMAP plots, velocity graphs are constructed using scv.tl.velocity_graph, and streamlines are plotted with scv.pl.velocity_embeding_stream. PAGA is run using latent velocities with scv.tl.paga, using the LatentVelo latent time as a prior. CellRank is used to compute terminal states. We combine the velocity kernel for LatentVelo’s velocities and the connectivity kernel using the nearest neighbour graph, with weights 0.2 and 0.8.

Slingshot^9^ trajectory analysis is performed to get detailed trajectories for specific cluster subsets^9^. Slingshot is run on 30 principle components, using the root clusters as informed by the LatentVelo analysis. To find lineage-associated genes, we use the tradeSeq^10^ function associationTest, and select genes with adjusted false discovery rate below 0.01 on each lineage separately. These genes are then used in trajectory heat maps to visualize the change in expression over pseudotime along the trajectory.

Palantir pseudotime^11^ is used to validate LatentVelo results on the epithelial subset. We choose a root at the GW10 budtip progenitors and run Palantir with 5 components. CellRank with a pseudotime kernel is used to compute terminal states and fate probabilities.

### Ligand-receptor interaction analysis

Cellchat^12^ was used to infer significant (p-val <0.01) ligand to receptor interactions between the fetal epithelia and stroma populations. Normalized counts and cell annotations from scRNAseq dataset split into early (GW 10-13), mid (GW 14-16), and late (GW 18-19) were used as an input. Source celltypes were subset according to spatial close celltypes to the target, as determined with Visium. Significant cell signaling pathways of interest were further analyzed. A paired Wilcoxon test for the sum of pathway interactions (information flow) is used to determine significantly varying pathways between early, mid, and late.

### In vitro- in vivo benchmarking of hPSC-derived fetal lung cells and organoids

CA1 (courtesy of Dr. Andras Nagy, Lunenfeld Tenanbaum Research Institute, Toronto) and BU3 hPSCs (courtesy of Dr. Darrell Kotton, Boston University) were used to generate hPSC-derived fetal lung cells and organoids using our previously established protocol^13^. Library preparation and single-cell analysis was done similarly as above. DEGs between gestational ages of the fetal lung epithelia was used to generate a panel of key features. These features were used as an input for pearson correlation analysis between hPSC-derived fetal lung cells and primary-derived fetal cells (grouped by GW). A combined embedding for the primary fetal lung epithelia and hPSC-derived cells was created by computing batch balanced nearest neighbors^14^ and computing a UMAP embedding. Clustering on this integrated space is done with Scanpy’s sc.tl.louvain with default resolution.

### Code availability

Code for the Trajectory analysis is available at https://github.com/Spencerfar/fetal_lung_velocity

### Immunofluorescence Staining

Previously archived formalin-fixed paraffin-embedded human fetal lung tissues were sectioned at 8um thickness for immunohistochemical analysis. Slides were deparaffinized using Xylene to 70% ethanol to water as previously described^28^. Briefly, antigen retrieval was then performed using either universal heat induced epitope retrieval (HIER, Abcam, Cat #ab208572) as per manufacturer’s protocol or citrate buffer solution (pH 6.0) for 10 minutes. The slides were then briefly rinsed with 1X PBS and blocked with 5% normal donkey serum and 0.5% BSA in PBS for 30 min at room temperature. In a humidified chamber, the slides were then incubated with primary antibodies (**Supplementary Table 4**) overnight at 4°C and then washed 2X with PBS. Secondary antibodies were then added for 1 hour at room temperature. Nuclei were counterstained with DAPI (ThermoFisher, 1:1000) for 15 minutes at room temperature and mounted with DAKO immunofluorescence mounting media and kept in the dark until imaging. Fluorescence images were captured with the Olympus spinning disc confocal microscope and analyzed with the Volocity imaging software (Quorum Technologies).

## Acknowledgement.

We would like to thank Zoe (Shuk-Yee) Ngan for her help in optimizing the single-cell library prep procedures. We would also like to thank Dien Nguyen for her help in data analysis. We would also like to thank Mr. Max Nilt and Ms. Dragica Curovic from the Research Centre for Women’s and Infants’ Health (RCWIH) BioBank for their help with the fetal lung tissue collection. We would also like to thank Mr. Alper Celik from the Centre of Computation Medicine (CCM) for his guidance in the bioinformatics analyses. Computational power was enabled by support provided by Compute Canada (www.computecanada.ca) and the internal SickKids High Power Computing (HPC) platform for processing our scRNA-seq datasets. Funding support for this work was awarded to Dr. Amy Wong by the SickKids Foundation & CIHR-IHDCYH grant (NI20-1070), and the Stem Cell Network Early Career Investigator - Innovation Award (FY21/ECI23). HQ is a recipient of the Data Science Institute Student Fellowship award (2022-2025) and the Ontario Graduate Studentship award (2020 and 2022). SF is supported by a University of Toronto Data Sciences Institute Postdoctoral Fellowship. KK is a recipient of the 2022 Canada Graduate Student Masters award. Schematic illustrations were created using BioRender.com.

## Author Contributions

HQ, KK, and MW performed experimental assays. HQ, SF, MW, PX, and PK performed computational analysis. HQ and APW designed the study, HQ, SF, MW, TJ, APW analyzed the data, and HQ, SF and APW wrote the manuscript. All funding support for this work was awarded to APW. All authors reviewed the data, read, and approved the manuscript.

